# High-resolution single-cell atlas of the human B cell compartment and immune microenvironment across tissues

**DOI:** 10.64898/2026.05.05.722858

**Authors:** Leire de Campos-Mata, Yong Sun, Likun Du, Hui Wan, Elisa Marani, Vera Nilsén, Takuya Sekine, Anne Marchalot, Christopher Tibbitt, Tobias Kammann, Elli Mouchtaridi, Demi Brownlie, Nicole Marquardt, Malin Flodström-Tullberg, Johan K. Sandberg, Matthias Lutolf, Lauriane Cabon, Jenny Mjösberg, Carl Jorns, Marcus Buggert, Lennart Hammarström, Xiaofei Ye, Qiang Pan-Hammarström

## Abstract

The immune system is a dynamic network of diverse cell types distributed across tissues. While mouse studies have highlighted the importance of tissue-localized B-cell responses in infection and tissue repair, research on human B cells has remained largely confined to peripheral blood. Here, we specifically investigated the human B-cell compartment and its interactions with the immune microenvironment across 10 tissues, using single-cell RNA sequencing and paired B-cell receptor sequencing. We mapped the full spectrum of B-cell populations and revealed diverse differentiation trajectories spanning naive, memory, and plasma cell types. Germinal center B cells and plasma cells showed tissue-specific adaptations in transcriptional states, isotype usage, and functional profiles. Plasma cell isotypes influenced both effector functions and predicted interactions with T cells. Finally, we defined tissue-specific residency gene modules that outperformed existing memory B-cell signatures. Together, this dataset serves as a foundation for systematically studying tissue-localized B cells and reveals how local microenvironments shape humoral immunity.

## Introduction

B lymphocytes are essential components of the adaptive immune system and are responsible for the production of antibodies that target and neutralize infectious agents. B-cell development initiates in the bone marrow, where specific genes of the immunoglobulin (Ig) loci are rearranged to express unique B cell receptors (BCRs) on the surface of individual B cells. This process, known as V(D)J recombination, ensures the establishment of a diverse BCR repertoire prior to antigen encounter. Immature B cells exit the bone marrow and migrate to secondary lymphoid organs to continue their development. Following antigen encounter, activated B cells either differentiate into short-lived plasma cells, give rise to early memory B cells^1^, or induce a unique genetic program to form specialized germinal centers (GCs) after cognate interaction with antigen-specific T cells^2^. GCs are transient structures where B cells undergo affinity maturation through somatic hypermutation (SHM) in the variable (V) regions of their Ig genes, followed by positive selection of B cells with increased affinity for their cognate antigen. Affinity maturation gives rise to a pool of antibody-secreting plasmablasts that commit to the plasma cell lineage or memory B cells that are poised to respond rapidly following a secondary encounter with antigen^3^. Antibody effector functions are further diversified upon antigen encounter through class-switch recombination (CSR), which is a DNA recombination process that replaces the Ig constant (C) region with a downstream C domain gene. Antibody affinity maturation and functional diversification, along with B cell differentiation, collectively shape the humoral and B cell-mediated responses.

Recent studies have reshaped our understanding of immunity by identifying tissue-resident immune cell populations that exhibit distinct functional properties and homing capacities compared to their circulating counterparts^4,5^. By remaining localized, cells of the innate immune system help control tissue inflammation and provide a frontline defense against infectious agents, whereas cells of the adaptive immune system establish tissue-localized immunological memory. The existence and maintenance of tissue-resident memory T (TRM) cells have been extensively studied for over two decades^6,7^. In contrast, the formal identification of tissue-resident memory B (BRM) cells was achieved only recently, through a study using a mouse model of pulmonary influenza infection^8^. In humans, B cells exhibit heterogeneous distribution patterns between circulating and tissue sites^9–12^. Despite ongoing efforts to create multi-tissue human immune cell atlases, markers that enable the identification of tissue-resident B cells are still lacking. Furthermore, the immune microenvironment, or niche, supporting tissue-resident B cells in humans remains poorly understood.

In this study, we constructed a comprehensive single-cell atlas of mature human B cells across ten donors and ten donor-matched tissues by integrating single-cell transcriptomics with paired BCR sequencing. This approach enabled high-resolution analysis of B-cell differentiation, isotype usage, and clonal relationships across lymphoid, mucosal, and peripheral tissues. We characterized both canonical and previously underexplored B cell states, revealing diverse memory and plasma cell differentiation trajectories with tissue-adapted transcriptional and functional profiles. We further uncovered site-specific GC dynamics, plasma cell specialization, and isotype-associated immunoregulatory features. Importantly, we identified tissue-specific transcriptional modules for MBC residency that outperformed published signatures based on mouse studies or TRM cells. Together, this dataset provides a foundational resource for the systematic study of human tissue-localized B cells and offers new insights into how local microenvironments shape humoral immunity.

## Results

### Multi-tissue single-cell profiling of the human immune compartments

To characterize the heterogeneity of immune cell types across the human body, we leveraged an established comprehensive tissue resource and standardized protocol for acquiring multiple tissues from deceased organ donors during organ procurement for clinical transplantation (the Immunology Human Organ Donor Programme, IHOPE^13,14^). The collected tissue samples included secondary lymphoid organs (spleen, lung-draining lymph nodes, and mesenteric lymph nodes), barrier tissues (ileum, colon, cecum, lung, and skin), as well as blood and liver (**Figures 1A and S1A**). We isolated single cells from matched tissue samples obtained from nine organ donors aged 26 to 84, and from one healthy blood donor (**Table S1**). Using this tissue resource, we performed single-cell RNA sequencing (scRNA-seq) paired with single-cell VDJ sequencing (scVDJ-seq) on B cells enriched through CD19 surface staining (up to 70%), supplemented with CD19^-^ cells from the same donors (**Figures 1B and S1B**).

**Figure 1.**
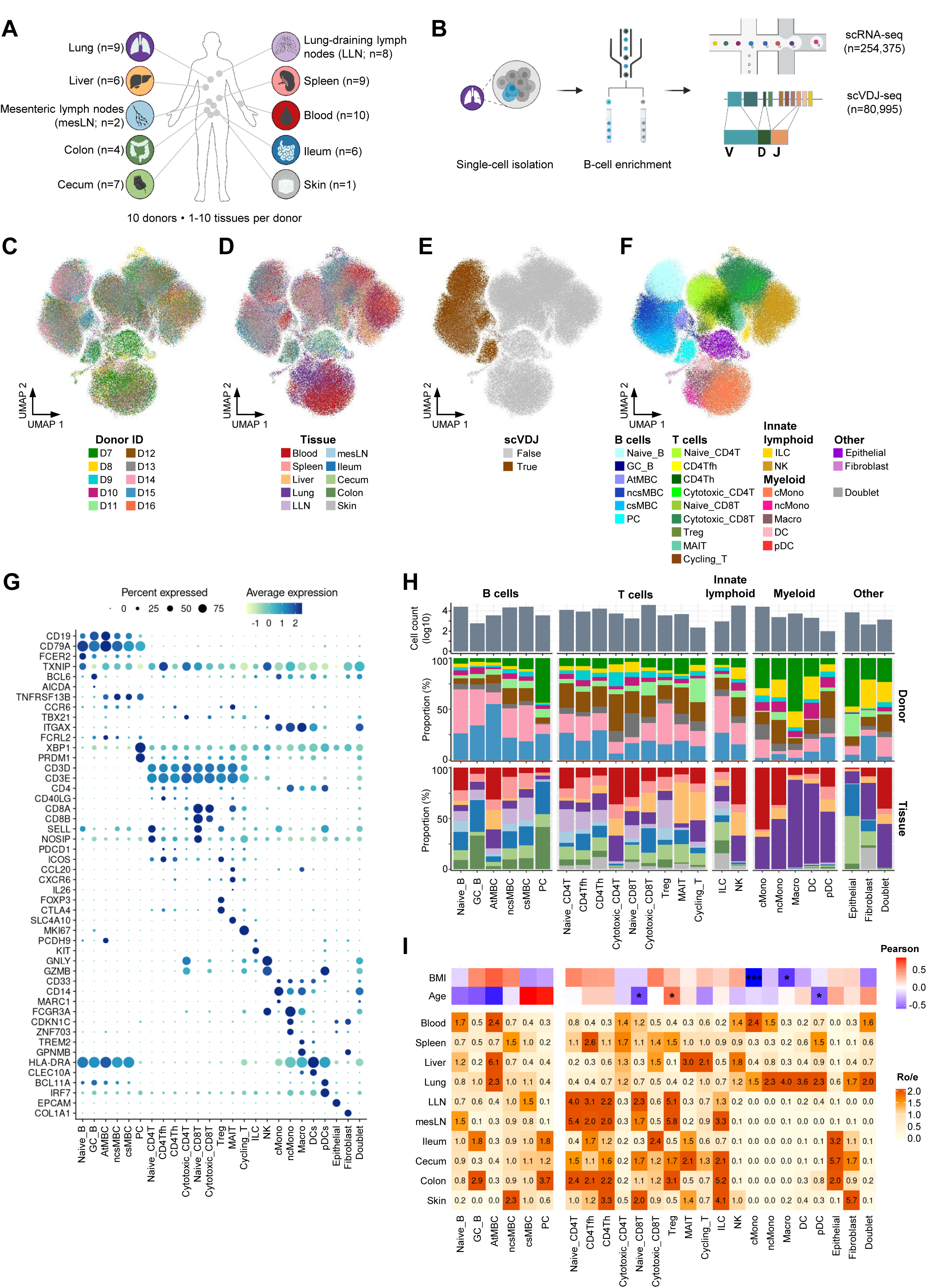
Single-cell profiling of the immune compartment across human tissues. (**A**) Schematic representation of the human tissue sample collection, including tissue name acronyms and the number of samples per tissue. (**B**) Overview of the experimental workflow for single-cell RNA sequencing (scRNA-seq) and single-cell VDJ sequencing (scVDJ-seq). (**C-F**) Uniform manifold approximation and projection (UMAP) plots of the immune cell compartment, colored by donor identity (**C**), tissue of origin (**D**), scVDJ data availability (**E**), and cell populations (**F**). (**G**) Dot plot of the average expression of marker genes for the identified cell populations. Dot size indicates the percentage of cells expressing each gene, and dot color represents the average gene expression level. (**H**) Bar plot showing the number of cells per cell population (upper panel), and stacked bar plots illustrating donor composition (middle panel) and tissue composition (lower panel) for each cell population. Donor identity and tissue origin are colored as in (**C**) and (**D**), respectively. (**I**) Upper panel: Heatmap of Pearson correlations between the donor body mass index (BMI) and age and the proportions of clusters within the B-cell and the non-B cell compartments calculated separately. Significance of correlations indicated by ***, p < 0.001; *, p < 0.05. Lower panel: Heatmap of tissue preference for each cluster within the B-cell and the non-B cell compartments separately, calculated by the Ratio of Observed to Expected (Ro/e) index.

Following stringent quality control, we retained the transcriptomes of 254,375 high-quality cells across 61 samples, encompassing 10 donors and up to 10 tissues ( **Figures S1C-S1E; Table S2**). The dataset was integrated across donors and tissues using Harmony^15^ to correct for batch effects, followed by unsupervised clustering based on gene expression. Uniform Manifold Approximation and Projection (UMAP) embeddings of the integrated dataset showed a homogeneous distribution of cells across donors and tissues of origin (**Figures 1C and 1D**). Cells were initially categorized into broad lineages based on known cell-lineage marker expression (**Figures S1F and S1G**) and scVDJ-seq data (**Figure 1E**). Next, we subgrouped these cells into 25 distinct cell populations (**Figures 1F and 1G**) with unique transcriptomic profiles (**Figure S1H; Table S3**).

Within the B-cell compartment, we identified 6 clusters representing the major B-cell sublineages of naive B cells (Naive_B), germinal center B cells (GC_B), memory B cells (AtMBC, ncsMBC, csMBC), and plasma cells (PC) (**Figures 1F-1H**). As expected, Naive_B cells were enriched in blood and secondary lymphoid organs, whereas PCs were highly enriched in the ileum and colon (**Figures 1H and 1I**). Although not statistically significant, the proportion of Naive_B, GC_B, atypical memory B cell (AtMBC), and non-class switched memory B cell (ncsMBC) clusters showed a negative correlation with donor age, while class switched memory B cells (csMBC) and PCs showed a positive correlation with age. A recent study showed that accumulation of Igs, especially IgG, correlated with age and senescence traits in mice^16^. In our dataset, the total proportion of IGHA-expressing cells, but not those expressing IGHG or IGHM/D, correlated positively with donor age (**Figures S1I-S1L**).

In the non-B cell compartment, naïve, helper, and regulatory CD4^+^ T cell, as well as naïve CD8^+^ T cell clusters, showed a strong enrichment in lung and mesenteric lymphoid organs (LLN and mesLN), whereas mucosal-associated invariant T (MAIT) cells were prevalent in liver and cecum. Myeloid cell clusters were highly enriched in blood and lung, and virtually absent from the gut and secondary lymphoid organs (**Figures 1H and 1I**). Epithelial cells were mainly enriched in gut tissues, whereas fibroblasts were enriched in different barrier tissues such as the lung, skin, and cecum. The proportion of naive CD8^+^ T cells (Naive_CD8T) and plasmacytoid dendritic cells (pDC) among non-B cells negatively correlated with donor age. In addition, the proportion of classical monocytes (cMono) showed a strong negative correlation with donor body mass index (BMI) (**Figure 1I**). Together, this integrated dataset constitutes a high-resolution, donor-matched atlas of immune cells across lymphoid, mucosal/barrier, and peripheral tissues, capturing the diversity of both B and non-B compartments and their variation with tissue context and age.

### Single-cell profiling of B-cell states

To improve resolution within the B-cell compartment, we reanalyzed 80,995 cells that expressed canonical B-cell markers (**Figure S1G; Table S4**). Unbiased clustering based on gene expression revealed 16 distinct B-cell clusters (**Figures 2A and 2B**), which we further characterized by BCR isotype usage and SHM frequency. SHM levels were categorized as “none” (SHM frequency = 0%), “intermediate” (SHM frequency < 20%), and “high” (SHM frequency > 20%) (**Figures 2C-2E and S2A-S2C**). To evaluate the robustness of our B-cell type annotations, we projected cell type labels from our B cell atlas onto an independent scRNA-seq dataset of human B cells (Dominguez Conde et al.^17^). The transferred labels aligned well with the cell type annotations of the external dataset (**Figure S2D-S2F**), supporting the reliability and biological relevance of our annotations. In addition, we validated the annotation of major B cell subtypes using gene set-based analysis by comparing our clusters with previously reported lineages from published datasets (**Figure S2G; Table S5**). Finally, although B-cell clusters displayed variation in donor and tissue composition, no clusters were uniquely associated with a specific donor or tissue of origin (**Figures S2H-S2K**).

**Figure 2.**
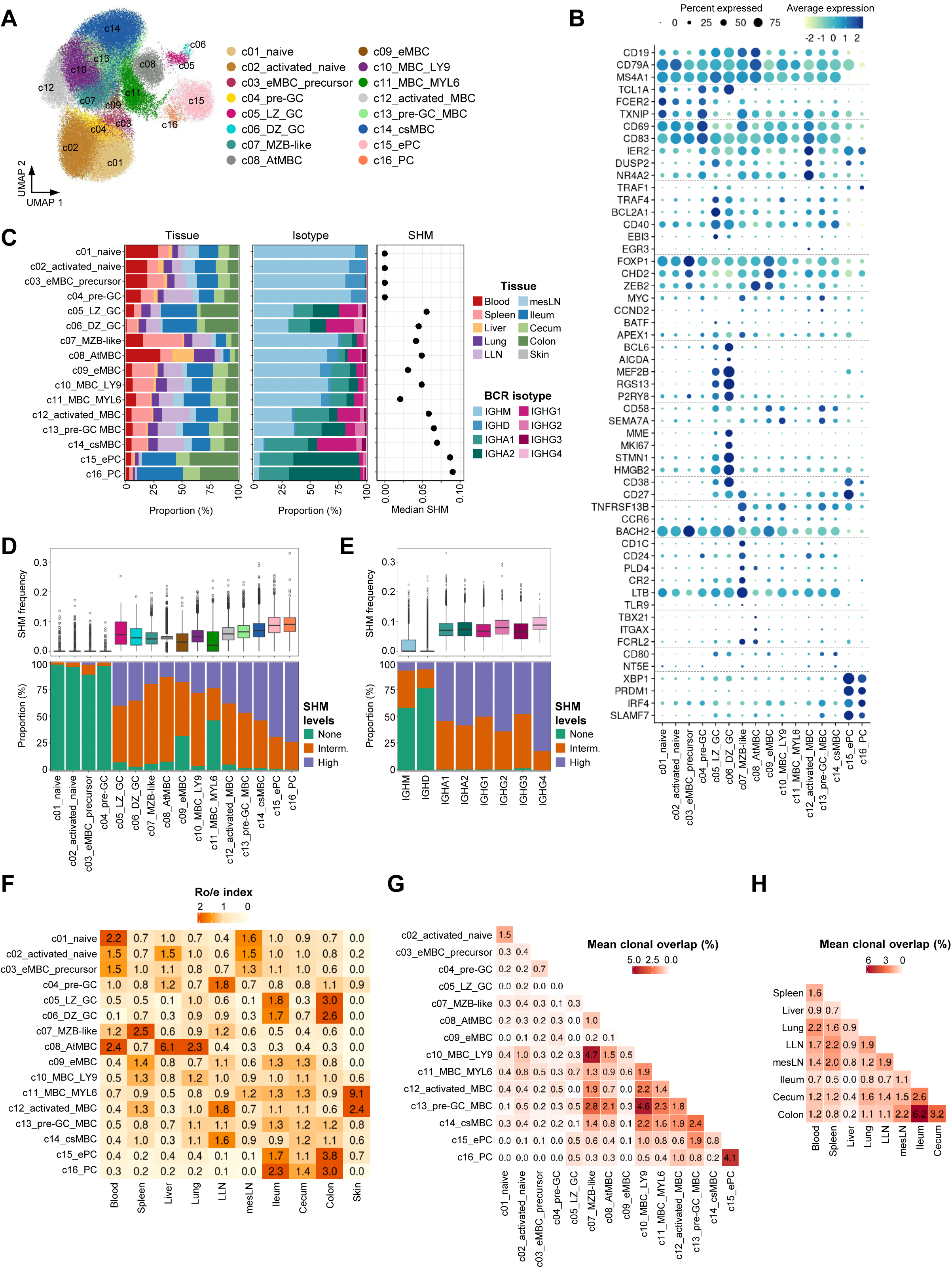
Single-cell profiling of the B-cell compartment. (**A**) UMAP plot of the B-cell compartment colored by identified B-cell clusters. (**B**) Dot plot of the average expression of marker genes for the identified B-cell clusters. Dot size indicates the percentage of cells expressing each gene, and dot color represents the average gene expression level. (**C**) Stacked bar plots showing the tissue composition (left panel), BCR isotype (center panel), and median somatic hypermutation (SHM, right panel) for each B-cell cluster. (**D**) Upper panel: Box plot of the SHM frequency on the IgH chain across B-cell clusters. Each box represents the interquartile range (IQR) of SHM frequencies for a given cluster, while the center line represents the median. Whiskers extend to 1.5 times the IQR starting from the respective box boundary. Individual dots represent outliers. Lower panel: Stacked bar plot showing the distribution of SHM levels across B-cell clusters. (**E**) Upper panel: Box plot of the SHM frequency on the IgH chain across isotypes. Each box represents the interquartile range (IQR) of SHM frequencies for a given isotype, while the center line represents the median. Whiskers extend to 1.5 times the IQR starting from the respective box boundary. Individual dots represent outliers. Lower panel: Stacked bar plot showing the distribution of SHM levels across isotypes. (**F**) Heatmap of tissue preference for each B-cell cluster calculated by the Ratio of Observed to Expected (Ro/e) index. (**G-H**) Heatmap of mean pairwise clonal overlap between B-cell clusters (**G**) or tissues (**H**) across donors. Values represent the average percentage of shared unique clones between cluster pairs, normalized by mean repertoire size. Clusters with fewer than 20 shared clones in the intra-cluster comparison were excluded.

In our dataset, we identified four naive B cell clusters (c01-c04), characterized by the expression of TCL1A, FCER2, and TXNIP, as well as IGHM and IGHD isotypes (**Figures 2B, 2C, and S2A**). As anticipated, these naive B-cell clusters exhibited no detectable SHM, distinguishing them from other B-cell clusters (**Figures 2C, 2D, S2B, and S2C**). Among the naive lineage, we identified three clusters of circulating naive B cells (c01_naive, c02_activated_naive, and c03_eMBC_precursor) enriched in blood and mesLN tissues (**Figures 2C and 2F**).

The c02_activated_naive cluster represents early activated naive B cells, characterized by moderate upregulation of early activation markers (*CD69*, *CD83*) and genes associated with the CD40 and BCR signaling pathways (*TRAF4*, *BCL2A1)* (**Figure 2B**). Differential expression analysis revealed high expression of *TMEM123* in c02_activated_naive, a gene linked to IL4-mediated B-cell activation ( **Figure S3A**). The c03_eMBC_precursor cluster showed high expression of genes that repress plasma cell differentiation, such as *FOXP1* and *BACH2* (**Figures 2B and S3A**), suggesting that this cluster may represent an early memory B cell (eMBC) precursor. Consistent with this notion, c03_eMBC_precursor showed increased expression of transcription factors (TFs) linked to non-GC-derived eMBC differentiation and inhibition of GC formation, including *CHD2* and *ZEB2*^18–20^ (**Figure 2B**). Additionally, we identified a pre-GC cluster (c04_pre-GC), representing a transitional state between activated naive B cells and GC B cells. Cells in this cluster expressed increasing levels of GC-associated markers, including *MYC*, *CCND2*, *BATF*, *BCL2A1*, and *BCL6*, while still retaining markers characteristic of naive B cells (**Figure 2B**). The c04_pre-GC cluster was moderately enriched for signatures of B cell activation^21^, lymphocyte costimulation, and KRAS signaling pathways, suggesting that these cells are primed for entry into the GC reaction (**Figure S3B)**. The positive correlations in cell abundances between c01_naive and c02_activated_naive, and between c02_activated_naive and c04_pre-GC, support a sequential activation trajectory leading to GC formation **(Figure S3C**).

GC B cells (c05-c06) expressed mostly class-switched, hypermutated isotypes (**Figures 2A-2D and S2A-S2C**), and were enriched in the ileum and colon (**Figure 2F**), presumably representing gut-associated lymphoid tissue (GALT). GC B cells engaged BCR and mechanistic target of rapamycin complex 1 (mTORC1) signalling pathways, as well as MYC pathways, all of which are essential for effective GC responses. Moreover, GC B cells showed antigen-presenting abilities, suggesting ongoing affinity selection within the GCs (**Figure S3B)**. The abundances of GC B cells correlated with c04_pre-GC, indicative of their frequent co-occurrence ( **Figure S3C**). Within the GC compartment, we identified two distinct clusters associated with light zone (LZ) and dark zone (DZ) phases: c05_LZ_GC and c06_DZ_GC, respectively. c05_LZ_GC cells showed upregulation of key GC-specific markers (ME2B, RGS13, BCL6, and AICDA), alongside markers characteristic of LZ cells (SEMA7A, CD58, EBI3, and BATF). In contrast, c06_DZ_GC B cells were distinguished by the high expression of proliferation markers (**Figures 2B and S3A**).

Marginal zone B (MZB) cells (c07) represent a unique subset of innate-like B cells which participate in rapid responses to blood-borne pathogens and secrete natural antibodies^22^. In our analysis, we identified a cluster of MZB-like cells (c07_MZB-like) that was highly enriched in the spleen. This cluster showed increased expression levels of *CD1C, CD24, PLD4, and CR2*, as well as LTB, the gene encoding the cytokine lymphotoxin beta (**Figures 2A-2F and S3A**). In mouse models, lymphotoxin beta is critical for the development of the splenic marginal zone, suggesting a similar role for c07_MZB-like cells in human tissue homeostasis. Additionally, B cells expressing lymphotoxin are known to support early innate immune responses in mouse models of infection^23^. Consistent with this, c07_MZB-like also expressed TNFRSF13B and TLR9, genes involved in early, T-cell independent B cell responses^24^ (**Figures 2B and S3A**). Despite predominantly expressing non-class-switched isotypes, c07_MZB-like cells accumulated a range of SHM levels, suggesting prior antigen exposure (**Figure 2D**). Clonal overlap analysis revealed that these cells shared clonal lineages with nearly all memory B cell clusters (**Figure 2G**). This broad connectivity suggests that c07_MZB-like cells may function as a founder population that contributes to the generation of diverse memory B cell fates. Altogether, these findings position c07_MZB-like cells as a versatile, antigen-experienced population with innate-like features and the potential to give rise to multiple branches of the memory B cell lineage.

B-cell clusters within the memory compartment were identified by the co-expression of TNFRSF13B and CD27, comprising subsets and states that differed both transcriptomically and in isotype usage. We identified a cluster, c08_AtMBCs, that displayed transcriptional features similar to those of atypical memory B cells (AtMBCs), which have previously been linked to infections and autoimmune diseases^25,26^ (**Figure S2G**). Specifically, c08_AtMBCs were highly enriched in blood, liver, and lung, and expressed high levels of *ITGAX, TBX21, and FCRL2*, alongside low expression of *CR2* and *CD27* (**Figures 2A-2F**). The abundance of the c08_AtMBC cluster did not correlate with donor age (**Figure S3C**). This cluster expressed mostly IGHM and IGHD isotypes with low SHM levels (**Figures 2C-2E and S2A-S2C**). Additionally, c08_AtMBCs expressed features of exhaustion and antigen presentation (**Figure S3B**). Notably, they expressed high levels of *ZEB2*, a transcription factor known to drive AtMBC differentiation and suppress *MEF2B*-mediated GC entry, reinforcing their proposed extrafollicular origin.

We further identified an early MBC cluster, c09_eMBC, distributed across multiple tissues (**Figures 2A-2C and 2F**). c09_eMBCs expressed high levels of CHD2, indicative of their early developmental nature. Based on transcriptional similarity, we hypothesize that c09_eMBCs arise from the c03_eMBC_precursor cluster (**Figure S3A**). However, this proposed relationship was not supported by clonal overlap analysis, as limited shared clones were detected between the two populations (**Figure 2G**). Like c08_AtMBCs, c09_eMBCs expressed ZEB2, supporting the notion of an extrafollicular origin.

Additionally, we identified an activated MBC cluster (c12_activated_MBC), which was enriched in spleen, LLN, and skin tissues. This cluster expressed hypermutated BCRs and markers of B-cell activation, yet showed no evidence of PC commitment (**Figures 2A-2F**). Moderate expression of GC-commitment markers, such as *MYC, CCND2, and BATF* (**Figure 2B**), suggests a potential for GC re-entry. Interestingly, the score for the B-cell activation signature correlated with the apoptosis signature (**Figure S3D**), pointing to a fine regulatory balance at the transcriptional level for the reactivation of MBCs. Comparative analysis with published datasets revealed that c12_activated_MBC shares transcriptional similarity with the c06_Bm_stress-response cluster from a pan-cancer B cell atlas ^27^, which has been associated with poor survival prognosis, as well as with the Bmemory-1 population of tumor-infiltrating B cells from hepatitis B virus (HBV)-associated diffuse large B-cell lymphoma ^28^ (**Figure S3E**). These associations suggest that c12_activated_MBC may represent a stress-responsive memory B cell state commonly found in tumor microenvironments. We also identified a cluster of pre-GC MBCs (c13_pre-GC_MBC), whose abundance significantly correlated with donor age (**Figure S3C**). This cluster expressed signatures of pre-GC stage and CSR, and markers of PC lineage commitment, such as *SLAMF7* and *IRF4* (**Figures 2B and 2G**). Finally, we identified a highly class-switched and somatically mutated MBC cluster (c14_csMBC) that expressed *CD80* and *NTE5* (**Figures 2A-2F**), markers associated with PC differentiation following secondary antigen exposure^29^. This cluster may preferentially differentiate into PCs upon local antigen re-encounter, rather than re-enter GCs for further affinity maturation.

In the PC compartment, we identified two distinct clusters (c15_ePC and c16_PC), both displaying a highly differentiated transcriptomic profiles relative to other B-cell clusters ( **Figure S3A**). PCs were enriched in the ileum and colon, and expressed heavily mutated *IGHA1* and *IGHA2* isotypes (**Figures 2A-2F**). The c15_ePC cluster displayed higher expression of major histocompatibility complex (MHC) class I genes compared to c16_PCs, suggesting that c15_ePCs may represent an intermediate stage preceding full PC maturation (**Figure S3B**). This early PC cluster also retained higher levels of B-cell lineage markers CD19 and CD79A, relative to mature c16_PCs. Differential expression analysis revealed upregulation of TXNDC5 in c15_ePCs, a gene previously linked to plasmablast fate programming^30^, further supporting its identity as an early PC population (**Figure S3A**). Functionally, c15_ePCs appeared more active, scoring highly for antibody secretion based on a gene signature specific to actively secreting B cells^31^ (**Figure S3B**). Clonal-overlap analysis showed that c15_ePCs shared multiple lineages with the c13_pre-GC_MBC population, while c16_PCs largely overlapped with c15_ePCs, supporting a trajectory in which GC-seeding MBCs first generate early plasmablasts that subsequently mature into tissue-localized PCs (**FigureB2G**). Consistent with extensive recirculation within the gut, total B-cell repertoires from the ileum and colon exhibited a high degree of clonal overlap (**FigureB2H**). Together, these findings outline a developmental continuum in which c15_ePCs act as transitional precursors that mature into the more specialized, terminally differentiated c16_PCs.

### Transcriptional features of B-cell activation and differentiation

Using transcriptome-based trajectory analysis and pseudotime inference, we mapped the dynamic progression of B-cell activation and differentiation across distinct lymphoid and nonlymphoid tissues (**Figures 3A and 3B**). At the start of the pseudotime trajectory, cells predominantly expressed *IGHM* and *IGHD* isotypes, reflecting the abundance of naive B-cell clusters ( **Figures 3B**, **S4A, and S4B**). As pseudotime advanced, these naive populations gradually declined, giving way to a wave of class-switched B cells primarily associated with MBC populations. Following this, a subsequent wave emerged, characterized by the predominance of *IGHA2*-expressing cells within the PC sublineage. The emergence of class-switched isotypes coincided in pseudotime with the accumulation of SHM and a gradual upregulation of the high-affinity signature score (**Figure 3C**). This progression effectively recapitulates the transition of B cells from a naive state to a high-affinity PC state.

**Figure 3.**
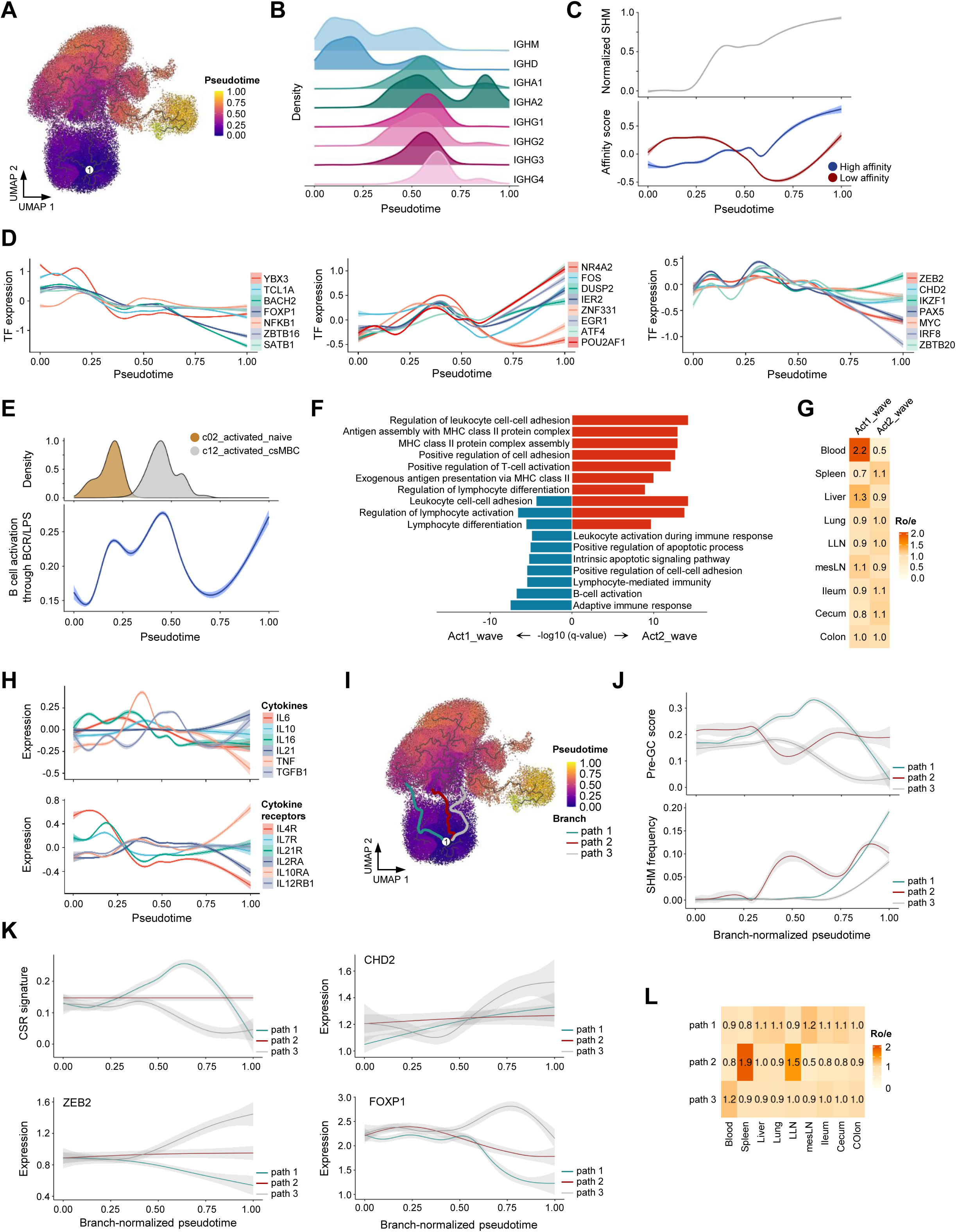
Trajectory analysis of B-cell activation and differentiation across immune tissues. (**A**) UMAP plot of the B-cell compartment colored by pseudotime. (**B**) Scaled density plots showing the relative distribution of B cells expressing different isotypes across pseudotime. (**C**) Two-dimensional plots showing the dynamic expression scores of normalized SHM and affinity gene signatures across pseudotime. (**D**) Two-dimensional plots showing the dynamic expression of selected TFs across pseudotime. (**E**) Scaled density plot showing the relative distribution of B-cell clusters c02_activated_naive and c12_activated_MBC across pseudotime (upper panel); and two-dimensional plot illustrating B-cell activation signature scores across pseudotime (lower panel). (**F**) Bar plot of the top Gene Ontology pathways enriched in cells from activation wave 1 (Act1_wave) and activation wave 2 (Act2_wave), as inferred from panel (E). Act1_wave cells were defined as those with B-cell activation scores > 0.2 and pseudotime between 0.15 and 0.27; Act2_wave cells were defined as those with activation scores > 0.2 and pseudotime between 0.35 and 0.55. (**G**) Heatmap of tissue preference for each activation wave calculated by the Ratio of Observed to Expected (Ro/e) index. (**H**) Two-dimensional plots showing the dynamic expression of selected cytokines and cytokine receptors along pseudotime. (**I**) UMAP plot of the B-cell compartment colored by pseudotime, highlighting three inferred developmental trajectories from naive to memory B cells: path 1 (turquoise), path 2 (red), and path 3 (light grey), overlaid on a gray trajectory skeleton. (**J, K**) Two-dimensional plots illustrating dynamic expression scores of pre-GC and CSR gene signatures, SHM frequency, and early extrafollicular memory B cell markers along pseudotime. (**L**) Heatmap of tissue preference for each developmental path calculated by the Ratio of Observed to Expected (Ro/e) index. (**C-E, H, J and K**). Solid lines represent linear fits, and shaded areas indicate 95% confidence intervals.

We performed DGE analysis to identify TFs upregulated at distinct stages along the pseudotime trajectory (**Figures 3D and S4C; Table S7**). Early in pseudotime, coinciding with the abundance of unswitched naive B-cell clusters, cells upregulated TFs such as YBX3, TCL1A, BACH2, and FOXP1. These TFs were progressively downregulated as pseudotime advanced, giving way to a second wave of upregulation, including TFs like NR4A2, FOS, DUSP2, and IER2. Interestingly, some of these TFs showed a biphasic expression pattern, being re-upregulated late in pseudotime, suggesting dual roles in MBC and PC differentiation. Additionally, we identified TFs that were upregulated during both the initial wave of unswitched isotype abundance and the subsequent wave of switched isotype abundance, highlighting their conserved roles in orchestrating B-cell maturation across different developmental stages. To determine whether specific TFs were associated with isotype-specific expression along the pseudotime trajectory, we divided pseudotime into three intervals: the first representing unswitched naive B cells, the second representing switched MBC and GC populations, and the third representing class-switched PCs (**Figures 3B, S4A, and S4B**). Within each interval, cells were further categorized by isotype, and all groups were systematically compared (**Table S8**). This approach revealed TFs uniquely upregulated under specific conditions (**Figure S4D**). While most TFs followed wave-specific patterns, some displayed condition-specific regulation. For instance, IGHG4-expressing cells during the second interval selectively downregulated *ZEB2* and *ZNF331*, highlighting subtle transcriptional regulation in specific isotype contexts. Additionally, NCOA3 was upregulated in *IGHG1*- and *IGHG2*-expressing cells within the third interval. As a known transcriptional target of XBP1, a key regulator of PC differentiation, *NCOA3* may act in concert with *XBP1* to facilitate IGHG-driven PC differentiation. Furthermore, *ZBTB38* was upregulated in *IGHA1*- and *IGHA2*-expressing cells during the third interval, suggesting a potential isotype-specific role in transcriptional control.

We next plotted the B-cell activation signature score along the pseudotime, which revealed two waves of B-cell activation ( **Figures 3E-3G; Table S9**). The first wave occurred early in pseudotime and overlapped with the peak abundance of the c02_activated_naive cluster. During this phase, B cells expressed genes involved in pathways of apoptosis, B-cell activation and differentiation, and cell-cell adhesion ( **Figure 3F**). Additionally, cells in the first wave of activation expressed higher levels of the cytokine gene IL16 and cytokine receptor genes IL4R, IL7R, and IL21R, underscoring the key role of these molecules in promoting a primary immune response (**Figure 3H**). As expected, this initial activation wave was followed by SHM accumulation and a transition from IGHM- and IGHD-expressing cells to those expressing class-switched isotypes (**Figures 3B and 3C**).

The second wave of activation aligned with the region where the c12_activated_MBC cluster was most abundant (**Figure 3E**). Cells in this second wave expressed pathways of T cell activation, cell adhesion, and antigen processing and presentation ( **Figure 3F**). This wave was characterized by upregulation of cytokine genes such as TNF, IL10, IL16, TGFB1, and a preceding peak in IL6 expression (**Figure 3H**). Additionally, the cytokine receptor profile shifted, with a downregulation of IL4R, IL7R, and IL21R and an upregulation of IL2RA, IL10RA, and IL12RB1, consistent with the role of activated MBCs in immune regulation and secondary immune responses. Following this wave, SHM accumulation and high-affinity scores^30^ further increased, culminating in the predominance of IGHA2-expressing PCs (**Figures 3B, 3C, and S4A-S4C**).

Interestingly, B cells from the initial activation wave exhibited a strong preference for peripheral blood, likely reflecting their migratory potential to home to their appropriate tissue niches (**Figure 3G**). In contrast, activated MBCs were predominantly localized at sites of inflammation and effector activity, with minimal circulation. Notably, cells in the second activation wave also showed a trend of preferential distribution in intestinal and respiratory tissues, suggesting their involvement in tissue-specific immune responses.

Pseudotime trajectory analysis further uncovered three distinct paths of naive-to-memory B-cell differentiation (**Figure 3I**). In path 1, the pre-GC signature score^32^ and SHM levels gradually increased, indicating a GC-dependent route of naive-to-memory differentiation. In contrast, path 3 exhibited low pre-GC signature and SHM levels, suggesting an extrafollicular pathway (**Figure 3J**). Supporting this notion, key TFs associated with early extrafollicular memory formation, including CHD2, ZEB2, and FOXP1, were specifically upregulated in path 3 (**Figure 3K**). Spatial preferences further distinguished the pathways. Path 1 showed a slight preference for mesLN, while path 3 favored blood (**Figure 3L**), consistent with its extrafollicular nature. Interestingly, path 2 was highly enriched in spleen and LLNs and exhibited expression of complement-related markers such as CR1, CR2, and CD1C, suggesting that path 2 represents a naive-to-MZB differentiation pathway (**Figure S4E; Table S10**). Together, these findings highlight the diverse routes of memory B-cell formation, each shaped by distinct transcriptional programs and tissue-specific contexts.

### Germinal center B-cell transcriptional and functional signatures across tissues

Although GC B cells were relatively scarce in our dataset, likely reflecting the healthy status of our adult donors with no ongoing inflammatory processes, we identified two distinct clusters of GC B cells (c05_LZ_GC and c06_DZ_GC) with distinct transcriptional programs ( **Figures S5A and S5B**). Given their low abundance, we analyzed GC B cells together across LLNs, ileum, and colon, the tissues with the highest GC representation (> 50 cells per tissue, **Figure 2C**). Of note, GC B cells were virtually absent from mesLNs. Despite the limited number of mesLN samples in our dataset (n = 2, **Figure 1A**), the low proportion of GC B cells in mesLNs (**Figures S2G and S2H**) suggests that, unlike in mice, human mesLNs may not serve as chronic inductive sites for mucosal immune responses.

GC B cells from LLNs showed enrichment for GO terms associated with respiratory burst and BCR signaling, suggesting robust antigen-driven activation and oxidative stress. In addition, LLN GC B cells expressed markers of exhaustion and high-affinity selection, with IGHG1 as the predominant BCR isotype. The exhaustion signature suggests a high antigenic load, likely due to persistent systemic antigen exposure in LLN GCs. In addition, LLN GCs expressed markers that might be relevant for the migration and retention of these cells in LLNs, such as integrins (ITGAE, ITGA4), CD22, and PTPN6 (involved in high endothelial venule-mediated trafficking), and H2ST1, a key enzyme in the biosynthesis of heparan sulfate, which facilitates chemokine-mediated lymphocyte homing to lymph nodes via endothelial cells **(Figures 4A-4F; Table S11**).

**Figure 4.**
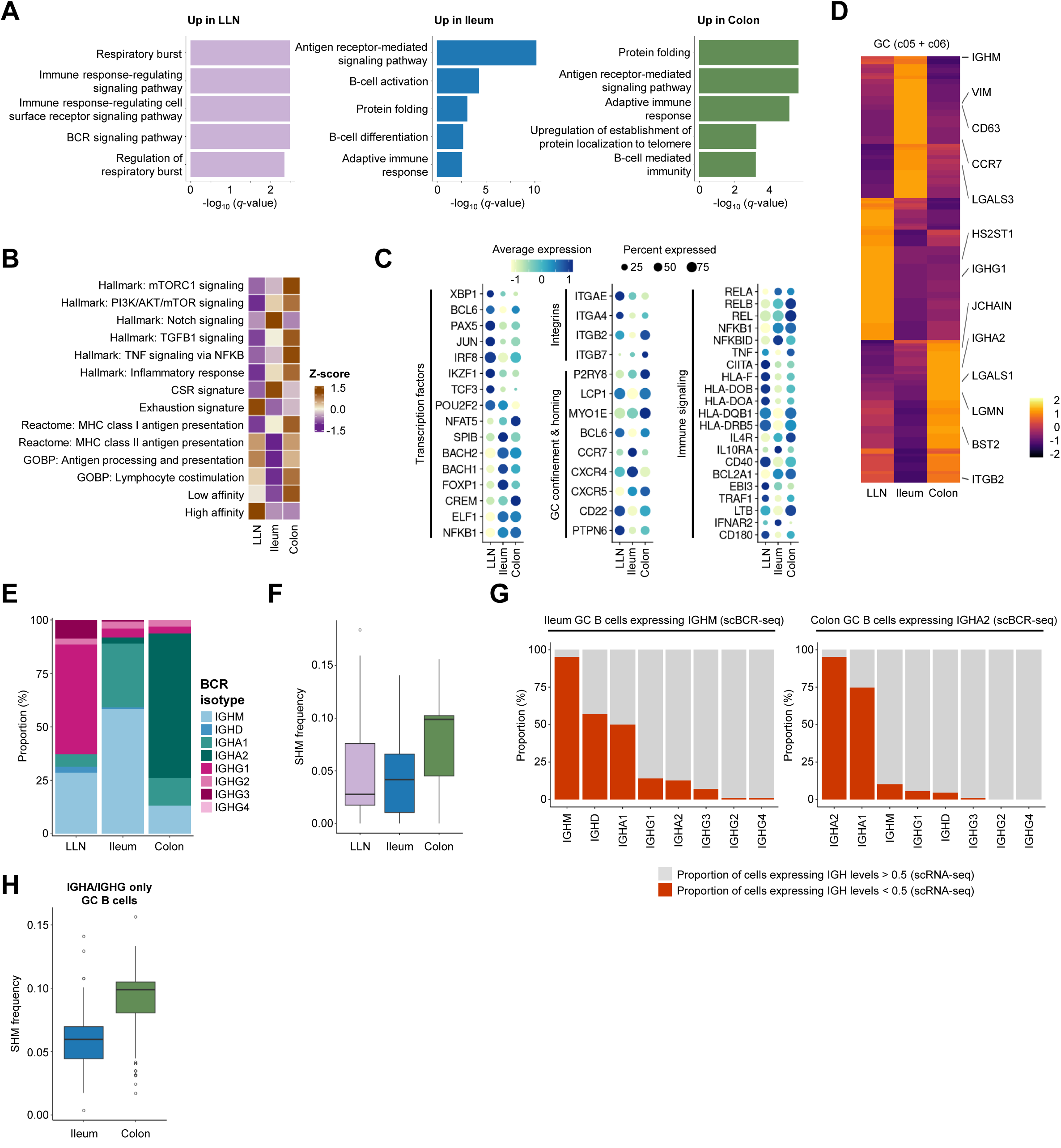
Tissue-specific transcriptional, functional, and isotypic adaptations of germinal center B cells. (A) Bar plots of top enriched Gene Ontology pathways in GC B cells by tissue; colors indicate tissue origin. (B) Heatmap showing average gene signature scores in GC B cells across tissues. (**C**) Dot plot of the average expression of marker genes for GC B cells across tissues. Dot size indicates the percentage of cells expressing each gene, and dot color represents the average gene expression level. (**D**) Heatmap showing the log_2_ fold change of the top 50 differentially expressed genes in GC B cells averaged across tissues. (**E**) Stacked bar plot showing BCR isotype composition in GC B cells by tissue. (**F**) Box plot of SHM frequency on the IgH chain in GC B cells across tissues. (**G**) Stacked bar plots showing the proportion of IGH gene-expressing cells based on scRNA-seq data. IGH expression levels are split by >0.5 (red) and ≤0.5 (gray). The analysis was performed separately for two GC B cell populations defined by scBCR-seq: ileal GC B cells expressing IGHM (left panel) and colonic GC B cells expressing IGHA2 (right panel). (**H**) Box plot of SHM frequency on the IgH chain of IGHA/IGHG-only GC B cells across tissues. (**F, H**) Each box represents the interquartile range (IQR) of SHM frequencies for a given cluster, while the center line represents the median. Whiskers extend to 1.5 times the IQR starting from the respective box boundary. Individual dots represent outliers. (**A-H**) Conditions with fewer than 50 cells were excluded from the analysis to ensure robust statistical power.

GC B cells from the ileum showed enrichment for GO terms associated with antigen-driven B-cell activation and maturation (**Figure 4A**). They also expressed signatures related to Notch signaling and CSR. In the ileum, GC B cells highly expressed *CCR7* and *CXCR4*, chemokine receptors critical for the homing of B cells to Peyer’s patches and the correct positioning within GCs. Additionally, ileal GC B cells expressed extracellular matrix and adhesion molecules (*VIM*, *ANXA2*, and *LGALS3*), markers characteristic of tissue-resident lymphocytes. Upregulation of *FOXP1*, *IL10RA,* and *IFNAR2* suggests a tissue-specific adaptation to handle inflammation and antiviral responses. The predominance of *IGHM* isotype in ileal GC B cells may reflect early-stage immune responses or a preference for innate-like humoral immunity in this environment. We detected *IGHA1* transcripts in *IGHM*-expressing GC B cells from the ileum (**Figure 4G**), although expression levels of *IGHA1* in single cells were insufficient to support the reconstruction of high-quality scVDJ contigs. This observation, together with a high CSR signature score^30^, may indicate ongoing CSR to IgA1 in ileal GC B cells.

GC B cells from the colon shared many GO terms with those from the ileum. Enrichment of functional signatures included activation of mTORC1, transforming growth factor beta (TGF-β) and nuclear factor kappa-light-chain-enhancer of activated B cells (NF-κB) signaling pathways, and an inflammatory signature, pointing to the induction of an inflammatory response, consistent with adaptation to the microbiota-rich and inflammatory environment of the colon. These cells exhibited a low-affinity, inflammatory response signature, with *IGHA2* identified as the primary BCR isotype. Thus, colon GC B cells appeared to be highly specialized for mucosal immunity, producing IgA2 to counter microbial antigens while tolerating a high inflammatory burden. The inflammatory signaling pathways and low-affinity maturation reflect the continuous antigenic stimulation within the colonic microenvironment with less rigorous selection. In addition, GC B cells in the colon upregulated markers of GC migration (*ITGB2*, *ITGB7*), confinement (*LCP1*, *P2RY8*, *MYO1E*, *BCL6*), and maintenance (*LTB* and *TNF*), which might help sustain GC reactions for longer. This could explain their higher rates of SHM in GC B cells from the colon compared to those from the LLN or ileum in cells expressing both class-switched and unswitched or only class-switched isotypes (**Figures 4F and 4H**). A high proportion of *IGHA2*-expressing GC B cells in the colon expressed coding *IGHA1* scRNA-seq (**Figure 4G**), which might point to an *IGHA1*-to-*IGHA2* sequential switching in the colon. In humans, the small intestine is dominated by IgA1 production, while the proportion of IgA2 becomes dominant in the distal colon^33^. Our results point to the local production of IgA1 and IgA2 in local lymphoid sites of the small and large intestine, respectively. Together, these findings highlight how tissue microenvironments shape GC B cell programs to meet distinct immune challenges.

### Tissue-imprinted transcriptional programs in human memory B cells

To dissect how tissue environments shape MBC states, we performed differential gene expression analysis across tissue sites. As anticipated, MBCs from the mesLNs, cecum, colon, and ileum exhibited similar transcriptional profiles. Additionally, MBCs from the blood and spleen clustered together and in close proximity to those from the LLNs, whereas MBCs from the liver and lung formed a distinct cluster (**Figure S6A; Table S12**). Analysis of Ig isotype expression revealed that blood-rich tissues, including blood, spleen, and liver, were enriched for IGHM-expressing MBCs with low levels of SHM. In contrast, MBCs from the respiratory tract (lung and LLN) showed an enrichment of IGHG1- and IGHG3-expressing cells. Meanwhile, gut-associated MBCs (mesLN, ileum, cecum, and colon) predominantly expressed IGHA1 and IGHA2 isotypes (**Figure S6B**).

Next, we assessed the expression of previously described residency-associated markers, largely derived from mouse studies and TRM literature. Tissue-resident cells typically downregulate lymphoid-homing markers while upregulating CD69, chemokine receptors, integrins, and adhesion molecules. In our dataset, expression of the lymphoid-homing receptor SELL, but not CCR7, was higher in blood and spleen MBCs compared to those from other tissues (**Figure S6C**). Instead, CCR7 expression was highest in the ileum, cecum, and colon, suggesting a role in MBC migration to GALT. CD69 expression was lowest in the lung and highest in the spleen, mesLN, and blood. The chemokine receptor CXCR3 was low across tissues but was increased in the lung and LLN, while CCR6 was more highly expressed in the spleen and lung compared to the blood. Additionally, CD44, a glycoprotein upregulated in murine lung tissue-resident memory B cells (BRM) and TRM, was expressed at lower levels in blood relative to other tissues. Gut-homing molecules such as CCR9, CCR10, GPR15, and ITGB7, were marginally expressed in our dataset.

To identify potential tissue-specific markers, we conducted a targeted analysis of genes encoding adhesion molecules, G protein-coupled receptors (GPCRs), and extracellular matrix components. Gene expression was compared across major tissue compartments, filtering for genes with expression >0.5 and at least a twofold difference between tissues (**Figure S6D**). An UpSet plot illustrated the intersection of gene sets across comparisons. As expected, SELL was enriched in MBCs from the blood (vs. gut and lung) and lymph nodes, consistent with its role in leukocyte trafficking. SCGB1A1 and RASGRP1 were consistently upregulated in the lung MBCs (vs. blood, gut, and LLN). *ICAM1*, *EXT1*, *GPR132*, *CXCR3*, and *PAG1* were upregulated in MBCs from the lung compared to those from the blood. *ICAM1* and *EXT1* enhance adhesion and extracellular matrix interactions, supporting tissue retention. *CXCR3* promotes migration to inflamed tissues, while *GPR132* and *PAG1* are implicated in immune regulation. *LGALS3* was elevated in MBCs from the gut (vs. blood, mesLN, and lung), and in the lung MBCs (vs. blood and LLN), suggesting a role in maintaining tissue-resident immune populations in both organs. *LGALS3* encodes galectin-3, a β-galactoside-binding protein involved in cell-cell adhesion and interactions with the extracellular matrix. Notably, galectin-3 has also been proposed as a marker of long-term T cell residency in human skin^34^, supporting its broader involvement in tissue-specific immune retention.

We next explored the potential role of LGALS3 in tissue residency by analyzing its top co-expressed genes in MBCs from the lung and gut compartments ( **Figure S6E**). In both tissues, ANXA2 and VIM, which are involved in cytoskeletal remodeling and cell adhesion, were highly correlated with LGALS3 expression, supporting their coordinated role in maintaining structural interactions within the tissue microenvironment. In the lung MBCs, LGALS3 was co-expressed with several S100 family members (including S100A6, S100A9, S100A10, and S100A11) and TXN, genes associated with stress response, inflammation, and epithelial function. In the gut MBCs, co-expressed genes included HMOX1, EMP3, and LMNA, which are involved in oxidative stress regulation, membrane dynamics, and nuclear architecture. LGALS3-expressing cells were broadly distributed across MBC clusters in both lung and gut, suggesting that galectin-3 expression reflects a tissue-adaptive state rather than a distinct MBC subtype. Notably, c12_activated_MBC showed consistently fewer LGALS3-expressing cells across tissues compared with other clusters (**Figure S6F**).

We validated the tissue-resident context of our dataset by comparing it with the dataset generated by Recaldin et al^35^, in which intestinal-immune organoids (IIO) were formed though the self-organization of epithelial organoids and autologous intestine-resident immune cells. We extracted B cells from their dataset (**Figures S7A and S7B**) and performed unbiased DGE analysis on B cells across the experimental conditions (**Figure S7C**; **Table S13**) and gene expression analysis based on custom gene lists with genes relevant to residency and migration (**Figure S7D; Table S14**). Next, we projected the expression patterns of genes upregulated in IIO B cells onto our MBC dataset, stratified by tissue of origin. Notably, genes highly expressed in IIO-derived B cells (top 50 genes: IIO; gene Cluster 3) showed the greatest transcriptional overlap with our gut-derived MBCs (**Figure S7E; Tables S13 and S14**), suggesting a shared residency-associated gene program and supporting a tissue-resident identity for intestinal B cells. In addition, LGALS3 and its co-expressed genes (ANXA2, VIM, and S100A11) were more highly expressed in both IIO-derived and tissue-resident B cells compared to circulating PBMC-derived B cells ( **Figure S7F**), further reinforcing their link to a residency-associated transcriptional program.

Markers such as CD69, while commonly used to identify tissue residency, are also upregulated upon B-cell activation. Similarly, chemokine receptors often associated with tissue-resident immune cells also mediate trafficking to lymphoid organs and GCs. To distinguish gene expression programs related to activation, differentiation, or tissue residency, we performed differential expression analysis of memory B cells across both tissue and cell type simultaneously. This analysis identified nine distinct gene expression modules, several of which reflected tissue-specific transcriptional programs (**Figures 5A and 5B; Table S15**). For example, Module_1 consisted exclusively of clusters from the LLN, while Module_2 grouped clusters from gut-associated tissues. Module_3 included clusters from blood, spleen, and LLN, consistent with a transcriptional program associated with circulating MBCs. Module_9 was composed of clusters from the lung, suggesting a lung-specific transcriptional signature. In contrast, Modules 6-8 grouped MBCs of the same cell type across multiple tissues, indicating that these cell populations maintain conserved transcriptional programs irrespective of their tissue localization.

**Figure 5.**
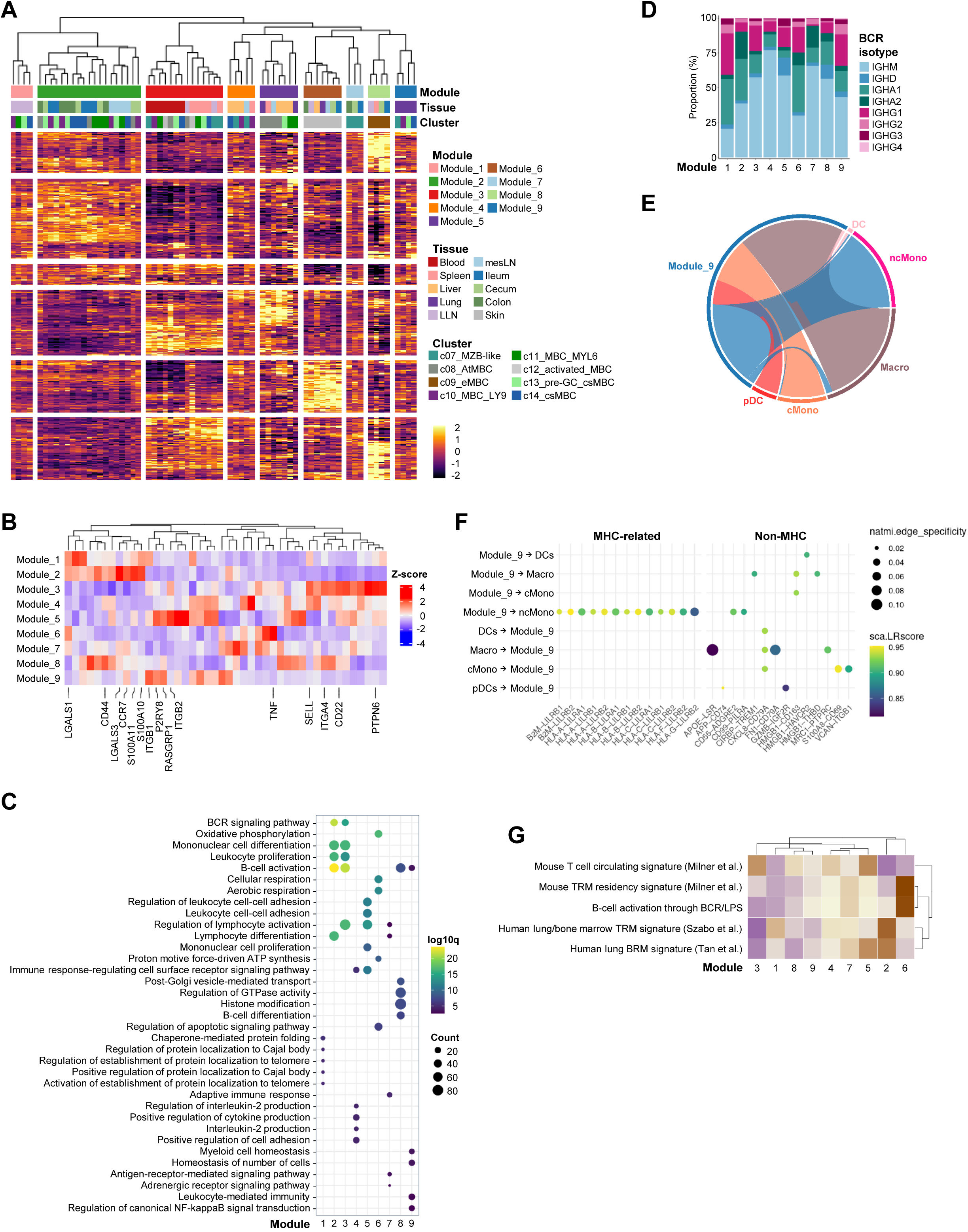
Tissue-specific transcriptional modules reveal functional specialization of memory B cells. (**A**) Heatmap showing the log fold change of the top 50 differentially expressed genes in MBCs stratified by cell type annotation and tissue of origin. Hierarchical clustering of columns revealed nine distinct transcriptional modules (Module_1 to Module_9). (**B**) Heatmap showing the average expression of selected genes among the top 50 differentially expressed genes in MBCs, stratified by module. Genes were chosen for their relevance to tissue residency or localization. (**C**) Dot plot showing GO enrichment analysis for each module. Dot color represents statistical significance (-log₁₀ adjusted *p*-value), and size represents gene count. (**D**) Stacked bar plot showing BCR isotype composition in MBCs by module. (**E**) Chord diagram showing predicted ligand-receptor interactions between Module_9 MBCs and myeloid cell subsets. (**F)** Dot plot showing predicted ligand-receptor interactions between Module_9 cells and myeloid cells, separated into MHC-related and non-MHC categories. Dot size represents interaction specificity scores, and color intensity reflects the average expression level of the ligand-receptor pair across the interacting cell types. (**G**) Heatmap showing average scores of previously published tissue-resident memory B cell (BRM) and T cell (TRM) gene signatures from human and mouse studies across the nine transcriptional modules.

We next performed GO enrichment analysis to define the functional specialization of each module (**Figure 5C**). Module_2 (gut-associated) was enriched for pathways involved in B cell differentiation and activation, likely reflecting continuous stimulation by the microbiota. Module_4 (liver-enriched) showed enrichment for cytokine production, consistent with an immunoregulatory role for hepatic B cells. Each module also displayed distinct Ig isotype usage (**Figure 5D**), Module_2 expressed IGHA1 and IGHA2, while respiratory-associated modules (Module_1 and Module_9) showed higher levels of IGHG1.

Because Module_9 was enriched for myeloid cell homeostasis pathways, we further investigated this lung-associated module using predicted ligand-receptor interactions (**Figures 5E and 5F**). These interactions linked Module_9 most strongly with non-classical monocytes (ncMono) and macrophages (Macro), followed by cMono and dendritic cell subsets. MHC-related signals (e.g., HLA–LILR) were largely specific to interactions with ncMono. Among non-MHC interactions, Module_9 B cells expressed *HMGB1*, targeting *CD163* on cMono, a potentially resident population. Other notable interactions included *CIRBP*–*TREM1* with Macro and *VCAN*–*ITGB1*, suggesting roles in inflammation and tissue remodeling.

To validate the tissue-specific transcriptional signatures identified in our clustering analysis, we assessed module-specific gene set scores in the Dominguez Conde et al. dataset, filtered for MBCs and age-associated B cells (ABCs), and stratified by organ (**Figures S2D and S2E**). Module_1 scored highest in MBCs from the thymus (THY), lamina propria of the jejunum (JEJLP), LLN, and mesenteric lymph nodes (MLN), consistent with its association with secondary lymphoid tissues. Module_2 was most enriched in gut-associated compartments, including JEJLP and the jejunal epithelium (JEJEPI), supporting its identification as a gut-specific signature. Module_3 showed the highest scores in spleen (SPL), bone marrow (BMA), THY, LLN, and blood (BLD), in line with a circulating or systemic MBC phenotype. Modules 4 and 5 were most prominent in the liver, while Module_9 scored highest in the lung (LNG) and omentum (OME) (**Figure S7G**). To further confirm the robustness of our gut-associated signature, we validated the enrichment of Module_2 in the Recaldin et al. dataset (**Figure S7H**).

Finally, we compared our transcriptional signatures with published BRM and TRM gene signatures from human and mouse studies^36–38^ (**Figure 5G**). When scored across our modules, these external signatures showed weak or inconsistent alignment with our tissue-specific clusters. In contrast, our module-derived signatures demonstrated strong, reproducible tissue associations across independent datasets. Together, these findings suggest that currently available tissue-residency markers may lack generalizability for human B cells and highlight the superior resolution of our approach in capturing the transcriptional imprint of tissue residency in MBCs.

### Region and isotype specific transcriptomic programs in plasma cell migration, maturation, and function

We identified two PC clusters representing sequential stages of differentiation. c15_ePCs expressed higher levels of MHC and B-cell lineage genes and were enriched for mitochondrial gene sets linked to active antibody secretion^31^ (**Figures S8A and S8B; Table S16**). In contrast, c16_PCs showed higher expression of extracellular matrix and survival genes, consistent with terminal differentiation^26^. To assess tissue-specific adaptations, we compared c15_ePC and c16_PC from blood, spleen, lung, LLN, cecum, ileum, and colon, the sites with highest PC abundances (> 50 cells per cluster and tissue, **Figure 2C**). PCs from blood-rich tissues displayed distinct transcriptomes from gut-derived PCs, indicating region-specific functional programs (**Figure 6A; Table S17**).

**Figure 6.**
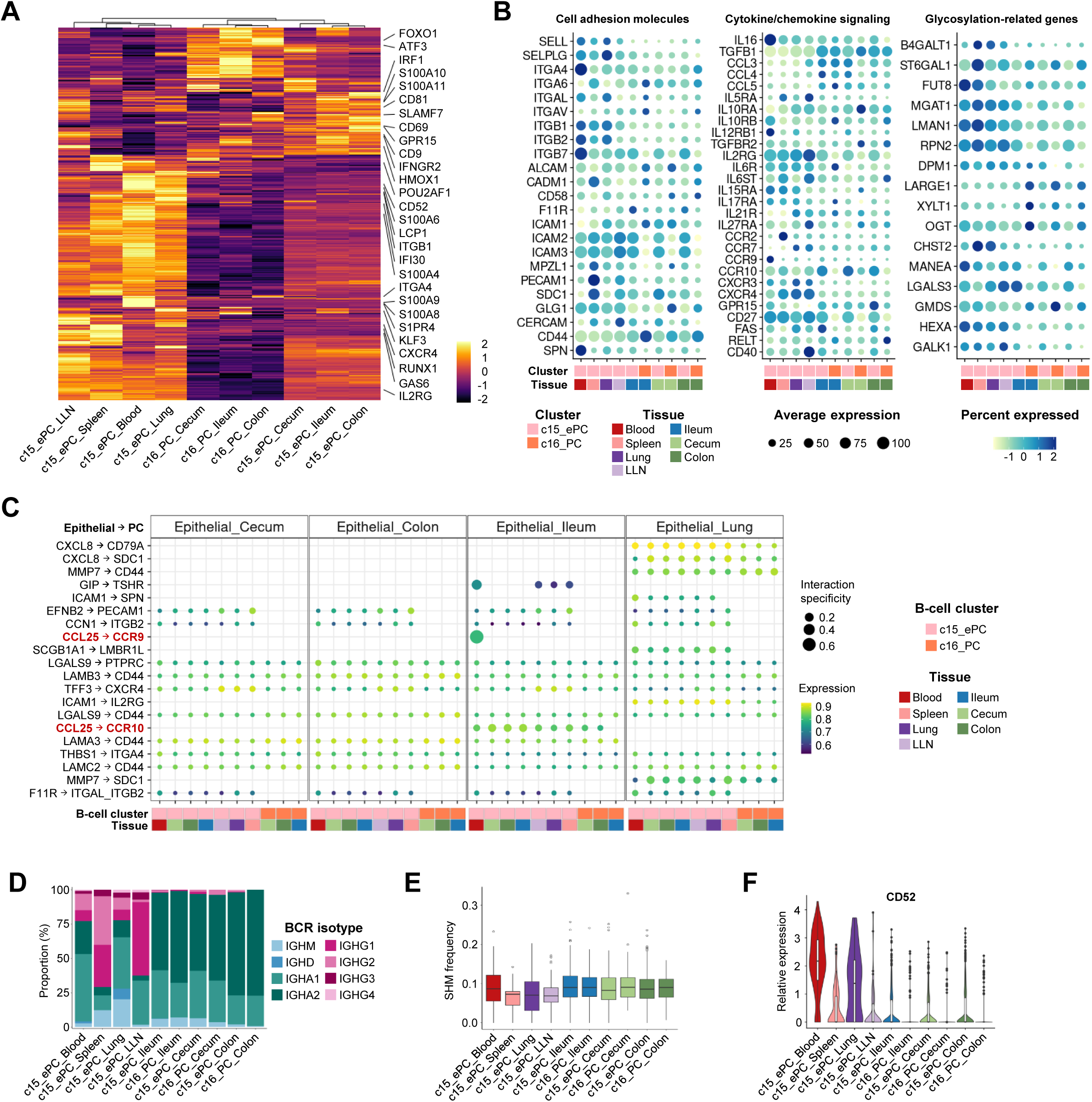
Tissue-resolved transcriptional adaptations of immature (c15_ePC) and mature (c16_PC) plasma cells. (**A**) Heatmap showing the log fold change of the top 50 differentially expressed genes in PCs averaged across tissues and cluster annotation. (**B**) Dot plot of the average expression of marker genes for the stated conditions. Dot size indicates the percentage of cells expressing each gene, and dot color represents the average gene expression level. (**C**) Dot plot depicting predicted ligand-receptor interactions between plasma cells and epithelial cells from different tissue origins. Dot size represents interaction specificity scores, and color intensity reflects the average expression level of the ligand-receptor pair across the interacting cell types. (**D**) Stacked bar plot showing BCR isotype composition in PCs by condition. (**E**) Box plot of SHM frequency on the IgH chain of PCs across conditions. Each box represents the interquartile range (IQR) of SHM frequencies for a given cluster, while the center line represents the median. Whiskers extend to 1.5 times the IQR starting from the respective box boundary. Individual dots represent outliers. (**F**) Violin plots showing the log-normalized expression of *CD52* across conditions. Each violin depicts the full expression distribution; the white box and central line indicate the interquartile range and median, respectively.

c15_ePCs from blood origin expressed S1PR4, a sphingosine-1-phosphate receptor that might point to the recent egress of these PCs from secondary lymphoid organs. These cells also expressed high levels of S100A8 and S100A9, which likely contribute to immune modulation during inflammatory responses. Additionally, circulating c15_ePCs expressed markers associated with tissue migration such as integrins (ITGA4, ITGB1, ITGB2, ITGB7) and chemokine receptors (CCR9, CXCR3, and GPR15) (**Figures 6A and 6B**). Transcriptomic analysis of PCs across isotypes highlighted upregulated ITGB1 in IGHG2-expressing PCs, suggesting a link between isotype expression and migratory behavior (**Figure S8A**). Cell-cell interaction analysis revealed potential interactions between circulating c15_ePCs and ileal epithelial cells, mediated by CCL25 expression in the epithelium and CCR9/CCR10 on PCs (**Figure 6C**). The migration of circulating c15_ePCs to the ileum is supported by the observation that approximately half of this cell population expressed IGHA1, the dominant isotype in the ileum (**Figure 6D**).

In the spleen, c15_ePCs expressed preferentially *IGHG1* and *IGHG2* isotypes (**Figure 6D**), reflecting the specialized role of the spleen in systemic immunity. These isotypes are critical for opsonization, complement activation, and effective neutralization of blood-borne pathogens. Additionally, splenic c15_ePCs showed high expression of genes involved in N-glycosylation, a process that fine-tunes antibody effector functions, including pathogen neutralization, complement activation and FCγ receptor binding. Splenic c15_ePCs also expressed a unique cell adhesion molecule profile, with higher expression of *CADM1*, *PECAM1*, and *MPZL1* (**Figure 6B**).

c15_ePCs from the lung displayed a heterogeneous distribution of antibody isotypes. Compared to LLN, lung PCs preferentially expressed IGHM, IGHA1, and IGHG2 isotypes, while IgGH1 and IGHG3 were more abundant in LLN c15_ePCs (**Figure 6D**). Lung c15_ePCs expressed high levels of selectins (SELL, SELPLG) and integrins (ITGAL, ITGB1, ITGB2), which likely facilitate their migration into inflamed tissues. Additionally, they expressed CHST2, an enzyme that sulfates glycosaminoglycans on adhesion molecules such as SELPLG and SELL, potentially enhancing their ability to adhere and traffic into inflamed or injured lung tissues (**Figures 6A and 6B**). Differential gene expression analysis highlighted tissue-specific differences in CD52 expression. PCs in the blood and lung expressed CD52 at higher levels compared to PCs from other tissues (**Figures 6A and 6F**). Additionally, transcriptomic analysis of PCs across isotypes showed CD52 upregulation in IGHM and IGHG2 isotypes (**Figure S8C; Table S18**). Clinical studies using alemtuzumab, an anti-CD52 antibody, resulted in the depletion of circulating T cells without eliminating a population of tissue-resident T cells in the skin^39^. Our findings suggest that differences in CD52 expression among PCs, likely influenced by their tissue of origin or isotype profile, may play a key role in determining their recirculating potential and capacity for tissue residency.

In the LLN, c15_ePCs expressed cytokine receptor genes involved in inflammatory signaling (IL27RA and IL6ST)^30^. The expression of IL5RA, typically linked to Th2 responses, suggests a role in allergic reactions and parasitic infections. Additionally, LLN c15_ePCS showed evidence of CD40-mediated interactions with T cells, which might support PC activation and survival. The expression of chemokine receptors CXCR3 and CXCR4 and the adhesion molecule CERCAM suggests the capacity of LLN c15_ePCs for chemotaxis toward inflamed or infected tissues (**Figures 6A and 6B**). Interaction predictions suggested that lung epithelial cells recruit PCs via the expression of CXCL8, a chemokine that promotes migration, MMP7, a metalloproteinase involved in tissue remodeling and invasion, and ICAM1, which supports leukocyte recruitment upon inflammation (**Figure 6C**).

PCs from the gut expressed only *IGHA1* and *IGHA2* isotypes and exhibited the highest levels of SHM compared to those from blood-rich tissues, although the differences were modest ( **Figures 6D and 6E**). c15_ePCs and c16_PCs of same tissue origin exhibited comparable SHM levels ( **Figure 6E**), supporting the idea that these populations represent distinct maturation states rather than deriving from separate precursors. Mature c16_PCs, but not c15_ePCs, from cecum and colon expressed high levels of *TGFBR2*, a receptor involved in TGF-β signaling, which is crucial for IgA class switching and mucosal immunity (**Figure 6B**). Mature gut c16_PCs expressed higher levels of glycosylation-related enzymes, including *LARGE1*, *XYLT1* and *OGT*, compared to immature c15_ePCs. This suggests that certain steps of glycosylation are enhanced during PC maturation in the gut, likely optimizing antibody secretion. In our dataset, ileum and cecum c16_PCs were responsive to IL-10 signaling, as evidenced by their expression of *IL10RB* and *IL10RA*, respectively, suggesting roles in immunoregulation and antibody production (**Figures 6A and 6B**). Immature gut c15_ePCs expressed *CCR10*, which in our dataset predicted the migration of PCs to the ileum. In addition, immature c15_ePCs from the colon expressed *GPR15*, which directs lymphocytes to the colonic mucosa (**Figures 6A and 6B**). Interestingly, mature c16_PCs from the gut lost cell-cell interactions with epithelial cells and downregulated the expression of homing receptors compared to their immature counterparts (**Figures 6B and 6C**). This downregulation of homing receptors underscores the transition of mature c16_PCs into a tissue-resident, non-recirculating state, reflecting functional specialization for sustained antibody production in the local mucosal environment.

B-cell isotype class can influence cell fate determination and effector function, which prompted us to investigate whether isotype-specific transcriptional programs shape PC function. To explore this, we analyzed GO terms associated with each isotype. *IGHG1*- and *IGHG2*-expressing PCs were enriched for GO terms related to oxidative phosphorylation, ATP synthesis, and mitochondrial processes, suggesting a metabolic profile tailored for energy-intensive antibody production. In contrast, IGHA2 and IGHM PCs expressed GO terms related to antigen presentation and MHC class II, indicative of roles in modulating adaptive immunity. Notably, IGHM PCs also expressed GO terms associated with regulation of T-cell activation, pointing to a potential immunoregulatory role (**Figure S8D**). IGHA1 PCs did not show statistically significant GO term enrichment. To further explore this immunoregulatory function, we examined interactions between isotype-specific PCs and T cells within the immunoregulatory network. IGHM- and IGHA-expressing PCs demonstrated unique interactions with T regulatory (Treg) cells via CCL3-CCR4 signaling, which is associated with recruitment and functional modulation of Tregs. This interaction was downregulated or absent in *IGHG1*- and *IGHG2*-expressing PCs. IGHG- and IGHA-expressing PCs, but not IgM-expressing PCs, interacted with T cells via CD86-CTLA4 interactions (**Figure S8F**). These findings suggest that isotype-specific transcriptional programs not only shape PC metabolic and immune profiles but also influence their capacity to engage with and modulate various T-cell subpopulations.

In summary, circulating c15_ePCs showed inflammatory and migratory traits, whereas splenic PCs specialized in systemic immunity, with increased glycosylation and cell adhesion. Lung and LLN PCs showed adaptation for migration and inflammatory responses, while gut PCs were focused on IgA production and maintaining mucosal tolerance. These results highlight tissue- and cell state-specific PC specialization that enables tailored immune responses.

## Discussion

This study provides the most comprehensive single-cell atlas of mature human B cells across lymphoid, mucosal, and peripheral tissues to date. By integrating transcriptomics, BCR analysis, and multi-tissue comparisons, we mapped the full spectrum of B-cell populations and uncovered diverse naive-to-memory differentiation pathways, including GC-dependent, extrafollicular, and marginal zone-like pathways. We also identified cytokine and TF expression dynamics that guided primary and recall B-cell activation. Our analysis of GC B cells revealed tissue-specific adaptations. LLN GCs expressed exhaustion and oxidative stress signatures; ileal GCs showed dominant IgM expression but ongoing CSR to IgA1; and colonic GCs were enriched for IgA2 and inflammatory markers. In parallel, PCs demonstrated both maturation-stage and tissue-specific specialization, ranging from migratory early plasmablast-like populations to long-lived resident effectors. We further found that PC isotype usage influences not only effector roles but also immunoregulatory interactions with T cells. We identified galectin-3 as a potential marker of memory B-cell residency in the lung and especially the gut. Finally, we identified tissue-specific transcriptional modules for MBC residency that outperformed published signatures based on mouse studies or TRM cells. Compared with previous datasets, our atlas provides higher annotation resolution, broader tissue coverage, improved transcriptomic and BCR integration, and deeper insights into the immune microenvironment that shapes B cell dynamics and development^17,40,41^.

Our study of naive B cells reveals how local tissue context and antigen exposure influence early differentiation pathways. The c03 naive cluster likely represents a flexible precursor population, supported by its clonal overlap with pre-GC (c04) cells and transcriptional similarity to early MBCs (c09). This suggests that c03 cells can commit to different fates, either progressing into the GC pathway or differentiating directly into MBCs. Consistent with this, mouse studies describe a precursor similar to c03 that gives rise to early MBCs like c09 under conditions of limited antigen^18^. The enrichment of c04_pre-GCs in LLNs, which actively support GCs, reflect naive B cells primed for GC entry. In contrast, the absence of c04_pre-GCs in mesLNs, which lack GCs and PCs and are instead dominated by naive and MBCs, emphasizes distinct immune landscapes in lymphoid tissues. Thus, antigen availability and tissue environment together may influence early B cell fate, with c04 enriched in LLNs primed for GC differentiation, while c03 and c09 predominate in gut and mesLNs, guiding alternative memory pathways.

Our data refine existing models of B-cell activation by revealing two sequential waves distinguished by location, function, and timing. The first wave involves circulating naive B cells that are primed toward a memory fate without committing to effector differentiation. This stage appears to be regulated by signals from Tfh cells and key TFs such as BACH2, which favor memory formation over GC or PC differentiation. Supporting this, ∼75% of GC B cells in our dataset were class-switched, suggesting that GCs are predominantly reseeded by reactivated MBCs rather than by naive precursors. The second wave consists of tissue-localized activated MBCs, which are absent from blood and show transcriptional programs linked to antigen presentation and T cell activation. Beyond local defense, these cells may help sustain or reinitiate GC reactions by presenting antigen to CD4^+^ T cells and amplifying Tfh responses. Our observation that activated MBCs accumulate in tissue sites and may reseed GCs is consistent with recent fate-mapping studies showing that local antigen re-exposure preferentially recruits tissue-resident primed B cells into recall GC responses, thereby enhancing affinity maturation and cross-reactivity at the site of initial priming^42^.

Our GC analysis highlights how mucosal environments shape B-cell maturation differently along the gut. Rather than functioning as isolated sites, ileal and colonic GCs appear to form a connected axis of IgA diversification. The CSR patterns and mutation levels are consistent with a model in which IgA1⁺ cells generated in ileal GCs migrate to the colon, where they undergo additional selection and further isotype refinement toward IgA2. The clonal connectivity between these sites supports this dynamic. Although GC B cells were relatively rare in our dataset, the moderate clonal overlap observed between GC B cells and PCs aligns with local production of IgA1 and IgA2-secreting PCs in the small and large intestine, respectively.

MBCs exhibit substantial transcriptional and functional diversity across tissues, reflecting adaptation to local immune microenvironments and supporting the presence of tissue-resident subsets analogous to TRMs. Certain MBC subsets, such as the LY9⁺ innate-like population enriched in the gut and lung, may be specialized for rapid, localized responses^43^, whereas others, like c09_eMBC, could represent terminally differentiated states with limited plasticity. Recent studies in mice and humans have reported long-lived MBCs in non-lymphoid sites with distinct phenotypes and functions^38,41,44^. In our dataset, conventional residency markers such as CD69, CCR6, and CXCR3 were variably expressed across tissues and did not consistently delineate residency. Instead, LGALS3 (galectin-3), co-expressed with cytoskeletal and epithelial-interacting genes, emerged as a candidate mediator of tissue residency, potentially anchoring MBCs in epithelial-rich niches. Its enrichment in lung- and gut-resident populations suggests a role in sustaining long-term local immunity, analogous to its function in skin TRMs^34^.

The tissue-specific transcriptional modules highlight how local immune contexts shape MBC specialization. In the lung, MBCs appear to be poised to interact with and regulate myeloid cells, supporting tissue integrity and local innate immunity. This is consistent with recent findings in mice showing that tissue-resident B cells spatially co-localize with and regulate macrophages, modulating their polarization and inflammatory status^45^. Gut MBCs seem adapted to chronic antigenic stimulation, likely reflecting ongoing engagement with the microbiota. Liver-enriched MBCs occupy a more effector-skewed niche, coexisting with cytotoxic T cells, MAIT cells, and NK cells within a tolerogenic environment, suggesting that local cellular composition and cytokine milieu shape their functional programs. Together, these patterns illustrate that tissue microenvironments strongly influence MBC localization, activation potential, and immune regulatory roles, emphasizing the need to consider niche-specific cues when studying MBC function or designing tissue-targeted immunotherapies.

Our analysis of PCs highlights how both maturation stage and tissue environment shape their identity. Although early and mature PCs represented distinct transcriptional states, their shared clonal origins indicate a continuum of differentiation from antigen-experienced B cells. The loss of CD19 and CD45 in mature PCs parallels features of long-lived intestinal PCs^4640^. In contrast, CD19-expressing PCs, like our c15_ePCs, align with a more transient, continuously replenished population^46^. Notably, PCs were more strongly stratified by tissue of origin than by developmental stage, emphasizing the dominant role of local cues in imprinting PC phenotypes^47^. The chemokine receptor landscape further supported this model. Early PCs displayed tissue-specific homing signatures, particularly gut-directed programs, while mature PCs showed diminished migratory potential consistent with long-term retention in situ. Isotype class added a second layer of specialization, with IgA⁺ and IgM⁺ PCs adopting transcriptional modules linked to antigen presentation and immunoregulation, in contrast to IgG⁺ PCs. These features align with emerging evidence that mucosal PCs participate not only in barrier defense but also in shaping local T cell responses, particularly through regulatory interactions in the gut^13,47^.

In addition to defining B-cell states in healthy tissues, our atlas provides a valuable reference for understanding B-cell phenotypes in disease. The AtMBC-like cluster we identified shares transcriptional programs with AtMBCs described in malaria, rheumatoid arthritis, and common variable immunodeficiency (CVID)^25,26^, suggesting that these disease-associated states have physiological counterparts that, when dysregulated, may contribute to pathology. Similarly, the activated MBC population resembles stress-responsive B cells reported in cancer and chronic viral infection ^27,28^, indicating that environmental stress in diseased tissues can drive chronic activation programs that are normally transient in healthy B cells. By anchoring these states within a high-resolution reference of human B-cell biology, our atlas provides a resource for dissecting how infections, autoimmunity, and cancer reshape B-cell differentiation trajectories. I t will also facilitate cross-study comparisons, enable annotation of scRNA-seq datasets generated in disease context, and aid in the identification of tissue-specific vulnerabilities or therapeutic targets.

In summary, this study provides a high-resolution atlas of human B-cell states across tissues, from naive and memory compartments to isotype-specialized plasma cells. It reveals several regulatory axes of B-cell differentiation, uncovers tissue-specific adaptation mechanisms, and introduces predictive transcriptional modules for tissue-resident subsets. These insights offer a valuable foundation for studying B-cell function in health and disease and hold promise for improving strategies in vaccination, mucosal immunology, and autoimmune therapy.

## Limitations of the study

This study has several limitations. First, the donor cohort is male-biased, limiting our ability to investigate sex-specific differences in B-cell phenotypes, particularly those unique to the female immune system. Second, although we profiled multiple lymphoid and non-lymphoid tissues, the donor and tissue diversity may not fully capture the complete range of B cell transcriptional programs across the human body. Third, cell abundance estimates may be affected by tissue-specific dissociation biases, potentially influencing the representation of certain B cell subsets. Finally, we did not perform spatially resolved transcriptomic analyses, which are required to contextualize B cell localization within tissue microenvironments. Despite these limitations, this work represents the most comprehensive B cell–focused single-cell atlas to date and provides a valuable resource for studying tissue-specific B cell biology.

## Material and methods

### Human sample collection and processing

Human organ tissues were obtained from deceased organ donors at the time of organ procurement for clinical transplantation within the IHOPE Programme. Criteria for inclusion were according to established national guidelines for organ donation and transplantation. The study was approved by the Swedish Ethical Review board (Dnr 2019-05016). Donors included in the study were free of cancer and seronegative for Hepatitis B, Hepatitis C, HIV, and syphilis. Hepatitis B serologic testing of donor D12 revealed a positive result for total antibody to hepatitis B core antigen (anti-HBc) but negative results for hepatitis B surface antigen (HBsAg) and Hepatitis B surface antibody (anti-HBs), indicating a resolved infection where anti-HBs levels have diminished. Donor demographics including age, sex, cause of death, and serological tests is summarized in **Table S1**. Organ donation was performed after declaration of brain death (DBD) or donation after circulatory death (DCD). Organs were then perfused *in situ* with cold organ preservation solution and maintained in cold preservation solution and transported to the laboratory within 2-4 hours of organ procurement. Tissue-processing protocols were adapted from protocols that have been previously described^13,14^. Briefly, mononuclear cells were isolated from the blood by standard density centrifugation using Lymphoprep (Axis-Shield) following the manufactureŕs instructions. Solid tissues were cut into small pieces and mechanically dissociated with the plunger of a syringe as a pestle. Half-dissociated tissues were then incubated in RPMI culture medium supplemented with 5 U/ml DNase I (Thermo Scientific) at 37°C for 1 hour at 120 rpm. Colon, cecum, and ileum samples were additionally complemented with 0.5 mg/ml Collagenase II and 0.5 mg/ml Collagenase IV (Gibco). After incubation, tissues were further mashed, filtered through a 70 µm nylon strainer, and washed with PBS. The number and viability of cells were evaluated. Single cells were then cryopreserved in fetal bovine serum (FBS) with 10% dimethyl sulphoxide (DMSO) and stored at -80°C before use.

### Fluorescence-activated cell-sorting (FACS)

Single cells were thawed in culture medium. Cells were blocked with human FcR Blocking Reagent (Miltenyi Biotec) for 10 min and stained with anti-CD19 BV421 in FACS buffer (PBS with 2% heat-inactivated FBS) for 20 min in ice. Right before sorting, cells were briefly incubated with propidium iodide (PI) staining solution. Live (PI-negative) CD19^-^ cells and live CD19^+^ cells were sorted in FACS buffer using BD FACSAria Fusion Sorter (BD Biosciences) using the strategy shown in **Figure S1A**.

### Single-cell gene expression and V(D)J library preparation and sequencing

Sorted live cells were enriched for CD19^+^ B cells using the strategy shown in **Figure S1A**. Using the Chromium Next GEM Single-Cell 5ü Reagent kit v2 (10x Genomics), live cells were loaded onto separate lanes of the Next GEM Chromium Controller (10x Genomics) for encapsulation (target recovery of 10,000 cells for each sample). Single-cell libraries were constructed using the manufacturer’s protocols. BCR sequencing libraries were prepared with Chromium Single Cell Human BCR Amplification Kit from 10x Genomics following the manufacturer’s protocols. Libraries were sequenced on a NovaSeq 6000 (Illumina) platform, aiming at 5,000 reads pairs per cell for the V(D)J library and 20,000 read pairs per cell for the 5’ Gene Expression library.

### scRNA-seq data quality control and integration

scRNA-seq data was aligned to the GRCh38 human reference genome and quantified using the Cell Ranger software (version 7.1.0, 10x Genomics). Putative doublets were identified and removed using Scrublet (0. 2. 3) ^48^. Cells with i) fewer than 200 or greater than 5,000 detected genes, ii) greater than 40,000 unique molecular identifier (UMI) counts, iii) greater than 20% mitochondrial reads, and iv) fewer than 5% ribosomal reads, were excluded. In addition, genes from the blacklist were removed as previously reported^49^, except for Ig and T cell receptor (TCR) genes. Downstream analysis of the filtered cell-gene matrix including data normalization and scaling was performed using Scanpy package (version 1.10.4) ^50^. Dimension reduction was performed using principal components analysis (PCA) based on the 3,000 most highly variable genes. Data integration across donors and tissues was performed using *Harmony* (version 0. 0. 9) ^51^. Visualization was performed using Uniform Manifold Approximation and Projection (UMAP).

### scRNA-seq data clustering and annotation

The harmony matrix was used for unsupervised clustering using the Leiden algorithm^52^. The clusters were assigned into five groups, including i) B cells, ii) T cells, iii) innate lymphoid cells, iv) myeloid cells, and iv) epithelial cells and fibroblasts. We removed cells that expressed mutually exclusive canonical markers and/or BCR genes. Eventually, a total of 254,375 high-quality cells were kept for further analysis. We performed Leiden algorithm-based subclustering for each group to increase the resolution of annotation. A total of 22 subclusters were generated and annotated using canonical cell type markers. In addition, a cluster of cells expressing mutually exclusive markers was identified and annotated as Doublet. The subclusters were then projected back to the UMAP of total cells. Additionally, paired scBCR-seq data were integrated with scRNA-seq data based on matched cell barcodes.

Using Leiden-based subclustering, we identified 16 distinct clusters within the B-cell compartment. To classify these clusters into specific cell types and states, we first assigned them to major B-cell lineages (naive B cells, GC B cells, MBCs, and PCs) by analyzing the expression of lineage-specific gene markers and literature-based gene signatures characteristic of these lineages. Further refinement of the annotations was achieved through manual curation, informed by an extensive review of the relevant literature.

### Label transfer to external dataset

To validate our B cell cluster annotations, we performed label transfer from our dataset to the single-cell RNA-seq dataset published by Dominguez Conde et al.^17^ using the Seurat v4 integration framework. Both datasets were log-normalized, and the top 3,000 highly variable features were identified using the FindVariableFeatures function. Anchors were computed between the reference (our dataset) and the query (Domínguez Conde et al.) using FindTransferAnchors, which applies canonical correlation analysis (CCA) to identify shared biological states, using the top 30 dimensions (dims = 1:30). Cell type labels from the reference were transferred to the query using TransferData, generating predicted identities for each cell in the query dataset. The predicted annotations were added to the Seurat object metadata and visualized with DimPlot to assess concordance between predicted labels and the structure of the query dataset.

### scBCR-seq analysis

scBCR-seq data was aligned and quantified using the cellranger-vdj software (version 7.1.0). Clonal identification was performed using the *Change-O (version 1. 3. 3)* repertoire clonal assignment toolkit and the SHazaM (version 1. 2. 0) R package^53^. First, the V gene, J gene, and junction were identified for each clone. Second, the productive clones were indicated and selected following the international immunogenetics information system (IMGT) definitions as i) having an open reading frame in the coding region; ii) having no defect in the start codon, splicing sites, or regulatory elements; and iii) having no stop codon and in-frame junction. Third, the clonotypes for the Ig sequences sharing the same V gene, J gene, and junction length were identified using nucleotide Hamming distance with automated threshold detection via the Gaussian Mixture Model. The Ig sequences of unclustered singleton clones were also identified as corresponding clonotypes. Finally, the germline Ig sequence for each donor was reconstructed to call the SHMs in each sample. The SHM level of each B cell was quantified as the point mutation rate in the IGHV sequence, and categorized as “none” (SHM frequency = 0%), “intermediate” (SHM frequency < 20%), and “high” (SHM frequency ≥ 20%).

### Tissue distribution preference analysis

To evaluate tissue distribution preferences of each B-cell cluster, we calculated the ratio of observed to expected cell numbers (Ro/e). The expected cell numbers were computed under the null hypothesis of random distribution across tissues using a chi-square test^27^. A Ro/e value greater than 1 indicates enrichment of a cluster in a specific tissue.

### Differential gene expression analysis

Differentially expressed genes (DEGs) between cell clusters were identified using the FindAllMarkers function of Seurat package with a Wilcoxon rank-sum test. On Figures 1 and 2, we removed BCR and TCR genes from the dataset before computing DEGs. For each cluster, differentially expressed genes were identified as the genes that (i) were detected in greater than 10% cells, (ii) had an absolute value of average log2 fold-changes greater than 1, and (iii) had an adjusted p-value less than 0.05. The top 50 DEGs per cluster, ranked by average log2 fold change, were used for visualization.

### Gene signature scoring analysis

The AddModuleScore function of Seurat was applied with default settings to calculate gene signature scores per cell. The mean signature score for each cluster was then calculated by averaging the scores across all cells within the cluster. Gene signature sets were collected from published studies and the Molecular Signatures Database (MSigDB)^54^. A detailed compilation of all sources for the gene signatures used is available in **Table S5**.

### Mean clonal overlap analysis

To quantify the mean clonal overlap between two repertoires we computed pairwise clonal persistence (CP) using a normalized metric. For any two repertoires, A and B, clonal persistence was calculated as:

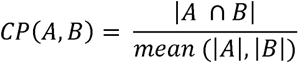

Where |A □ B*∣* denotes the number of unique clones shared between repertoires A and B, and *∣*A*∣* and *∣*B*∣* are the numbers of unique clones in A and B, respectively. The value was reported as a percentage.

### Gene ontology enrichment analysis

To identify enriched biological processes associated with DEGs across conditions, Gene Ontology (GO) term enrichment analysis was performed using clusterProfiler v4.8.3. For each condition, DEGs with an adjusted p-value < 0.05 were selected and mapped to ENTREZ gene IDs and the org.Hs.eg.db annotation database. GO enrichment analysis was conducted using the enrichGO function, focusing on the Biological Process (BP) ontology. The analysis used the Benjamini-Hochberg (BH) method for multiple testing correction, with a significance threshold of adjusted p-value < 0.05. Minimum and maximum gene set sizes were set to 10 and 500, respectively. Results were simplified using the simplify function to reduce redundancy in GO terms, and the top 5 or 10 terms per condition were selected for visualization.

### Pseudotime analysis

Pseudotime analysis of B cell populations was performed using the Monocle3 package (v1.3.4) to reconstruct differentiation trajectories. A Seurat object was converted into a CellDataSet, with cluster identities and UMAP embeddings transferred to enable trajectory inference. The learn_graph function was used to fit a principal graph onto the UMAP, represented as “skeleton lines” that outline the differentiation trajectory. Based on prior biological knowledge, c01_naive B cell cluster was selected as the root of the trajectory. Cells were then ordered along the trajectory using the order_cells function, which calculated pseudotime values for each cell. Pseudotime values were normalized and divided into 20 bins to study the BCR isotype, tissue and cluster composition per each bin.

Trajectory analysis revealed three major divergent pathways of naive-to-memory differentiation (path 1, path 2, and path 3). Cells belonging to these pathways were subsetted using the choose_graph_segments function, and pseudotime values for each path were normalized to allow for direct comparisons between trajectories.

### Cell-cell interaction analysis

To examine the expression of ligand-receptor pairs between different cell clusters, we used LIANA package^55^. Cell clusters with fewer than 50 cells were excluded from the analysis. A subset of cells representing the specified clusters was analyzed using the liana_wrap function, which integrates multiple ligand-receptor inference methods to predict communication between cell groups. The default OmniPath intercellular interaction resource was used as the ligand-receptor reference. Interactions were aggregated using the liana_aggregate function to rank ligand-receptor pairs based on their significance and consistency across methods.

### Statistics

Statistical analysis was performed using the Mann–Whitney U test. Significance was determined at p<0.05. Multiple test corrections were performed using false discovery rate (FDR).

## Acknowledgements

This work was supported by the Swedish Research Council (QPH), Swedish Cancer Society (QPH), the Knut and Alice Wallenberg Foundation (KAW2020.0101, LH and QPH), and the KAW Scholar Grant (QPH). IHOPE project was funded by the KAW Foundation (KAW2022.0021). Carl Jorns was supported by Region Stockholm (clinical research appointment, ALF).

