## Supplemental Figures for "High-resolution single-cell atlas of the human B cell compartment and immune microenvironment across tissues"

### Figure S1, related to Figure 1

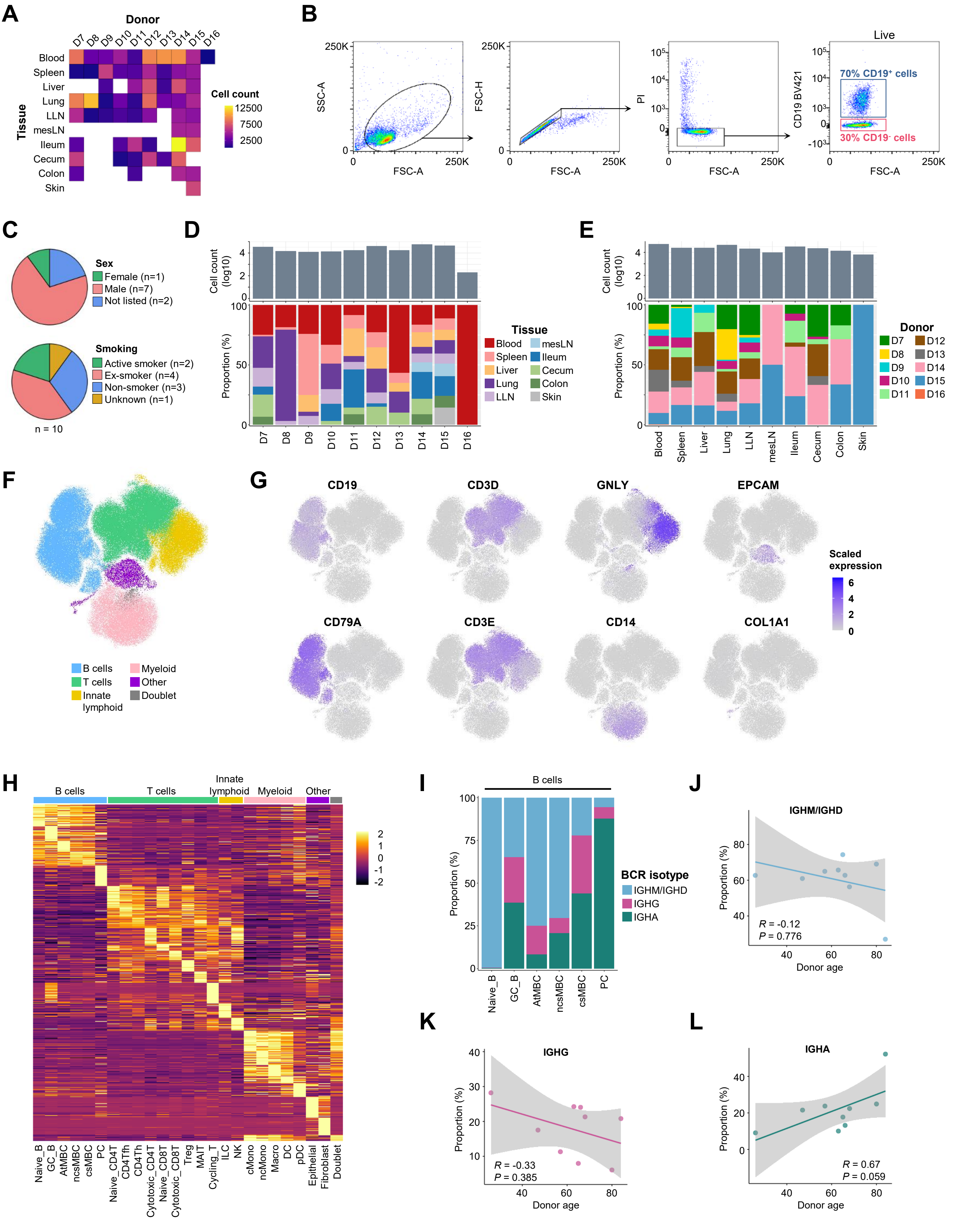

**Figure S1. Overview of the dataset, related to Figure 1.** (**A**) Tile plot displaying the distribution of donor samples across tissues. Tiles are colored by cell count, with only tiles corresponding to available samples shown. (**B**) Gating strategy used for FACSorting of live $PI^{-}CD19^{+}$ and live $PI^{-}CD19^{-}$ cells for scRNA-seq and scVDJ-seq. (**C**) Pie charts of donor demographics shown as reported sex (Female, n = 1; Male, n = 7; Not listed, n = 2) and smoking status (Active smoker, n = 2; Ex-smoker, n = 4, Non-smoker, n = 3, Unknown, n = 1). (**D**) Bar plot showing the number of cells per donor (upper panel), and stacked bar plot illustrating tissue composition (lower panel) for each donor. (**E**) Bar plot showing the number of cells per tissue (upper panel), and stacked bar plot illustrating donor composition (lower panel) for each tissue. (**F**) UMAP plot of the immune compartment colored by main cell identity. (**G**) UMAP plots of the immune compartment colored by the expression of cell-lineage marker genes for B cells, T cells, innate lymphoid cells, myeloid cells, epithelial cells and fibroblasts. (**H**) Heatmap showing the $\log_2$ fold change of the top 50 differentially expressed genes, averaged per cluster. (**I**) Stacked bar plot showing BCR isotype composition for each B-cell cluster. (**J-L**) Pearson correlation of BCR isotype proportions and donor age for *IGHM*/*IGHD* (**J**), *IGHG* (**K**), and *IGHA* (**L**).

### Figure S2, related to Figure 2

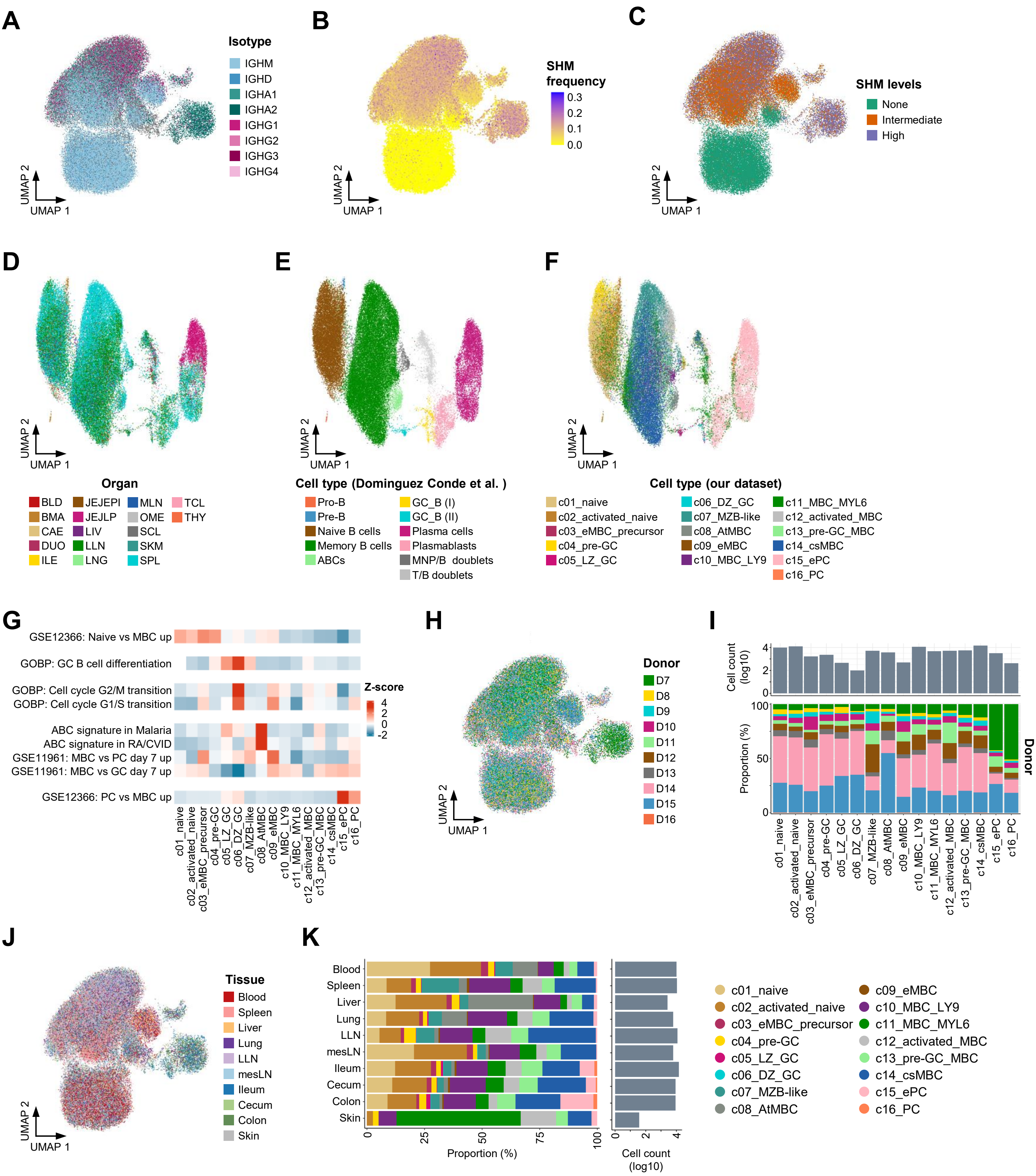

**Figure S2. Overview of the B-cell compartment, related to Figure 2.** (**A**) UMAP plot of the B-cell compartment colored by identified BCR isotype. (**B**) UMAP plot of the B-cell compartment colored by identified somatic hypermutation (SHM) frequency. (**C**) UMAP plot of the B-cell compartment colored by SHM levels. (**D**) UMAP plot of the Dominguez Conde et al. dataset, coloured by organ as annotated in the original publication. (**E**) UMAP plot of the Dominguez Conde et al. dataset, coloured by cell type as annotated in the original publication. (**F**) UMAP plot of the Dominguez Conde et al. dataset, colored by cell type labels transferred from our dataset using Seurat label transfer. (**G**) Heatmap showing average gene signature scores for each B-cell cluster. (**H**) UMAP plot of the immune compartment colored by donor identity. (**I**) Bar plot showing the number of cells per B-cell cluster (upper panel), and stacked bar plot illustrating donor composition (lower panel) for each B-cell cluster. Donor identity is colored as in (**H**). (**J**) UMAP plot of the B-cell compartment colored by tissue origin. (**K**) Stacked bar plot illustrating cluster composition for each B-cell cluster (left panel), and bar plot showing the number of cells per B-cell cluster (right panel). BLD, blood; BMA, bone marrow; CAE, caecum; DUO, duodenum; ILE, ileum; JEJEPI, jejunum (epithelial fraction); JEJLP, jejunum (lamina propria fraction); LIV, liver; LLN, lung-draining lymph nodes; LNG, lung; MLN, mesenteric lymph nodes; OME, omentum; SCL, sigmoid colon; SKM, skeletal muscle; SPL, spleen; TCL, transverse colon; THY, thymus.

### Figure S3, related to Figure 2

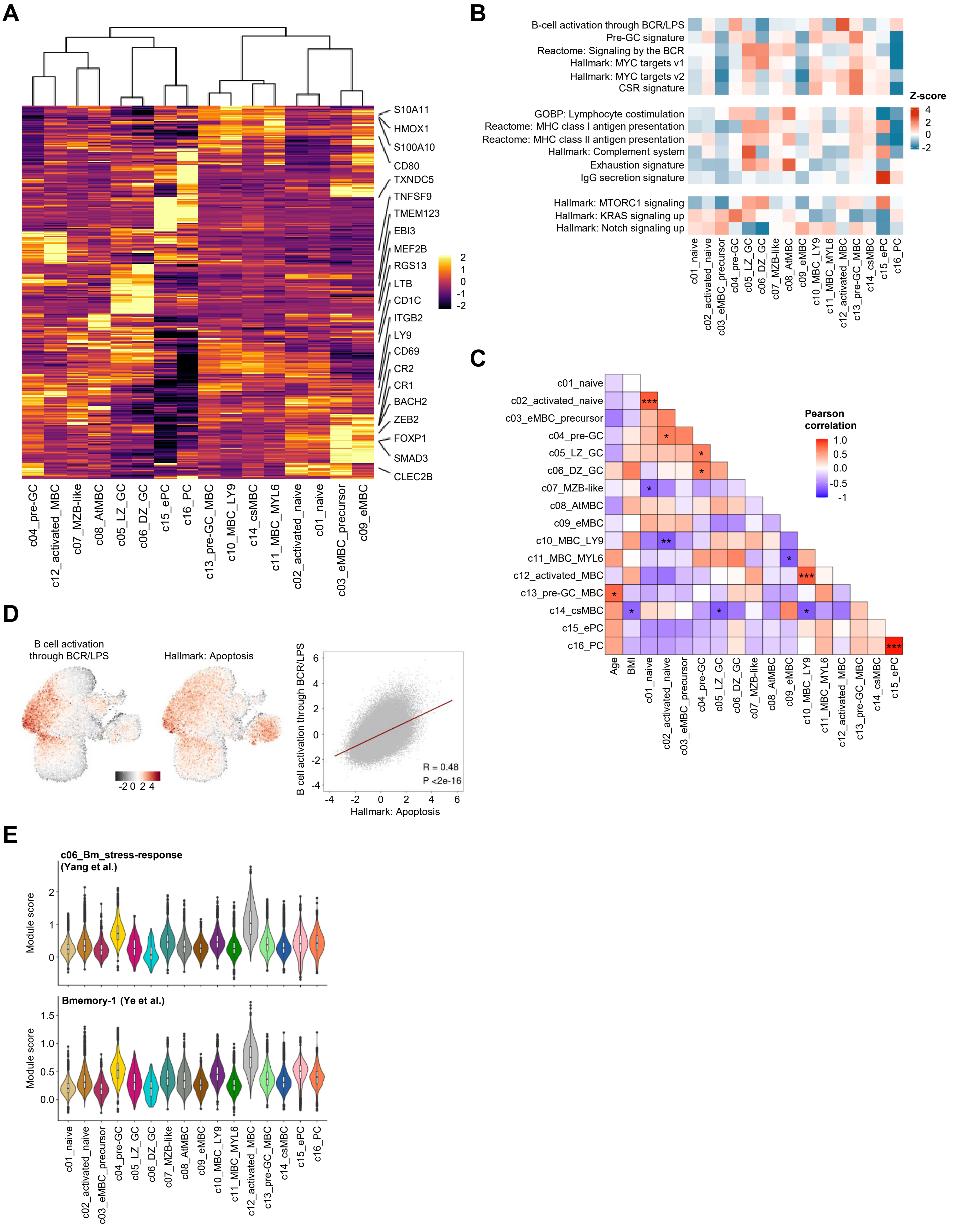

**Figure S3. Heterogeneity of B-cell clusters in the dataset, related to Figure 2.** (**A**) Heatmap showing the log fold change of the top 50 differentially expressed genes, averaged per B-cell cluster. A hierarchical clustering dendrogram shows the relationships among B-cell clusters based on gene expression profiles. (**B**) Heatmap showing average gene signature scores for each B-cell cluster. (**C**) Heatmap of Pearson correlation coefficients between B-cell cluster proportions, donor age, and body mass index (BMI). Cluster proportions were normalized within each donor and later tested for Pearson correlation. Significance of correlations indicated by ***, $p < 0.001$; **, $p < 0.01$; *, $p < 0.05$. (**D**) UMAP plots of the B-cell compartment colored by the indicated gene signature scores, as well as scatter plot of the Pearson correlation between the indicated gene signature scores across cells. (**E**) Violin plots showing the average module scores for the c06_Bm_stress-response signature from Yang et al. (upper plot) and the Bmemory-1 signature from Ye et al. (bottom panel) across B-cell clusters.

### Figure S4, related to Figure 3

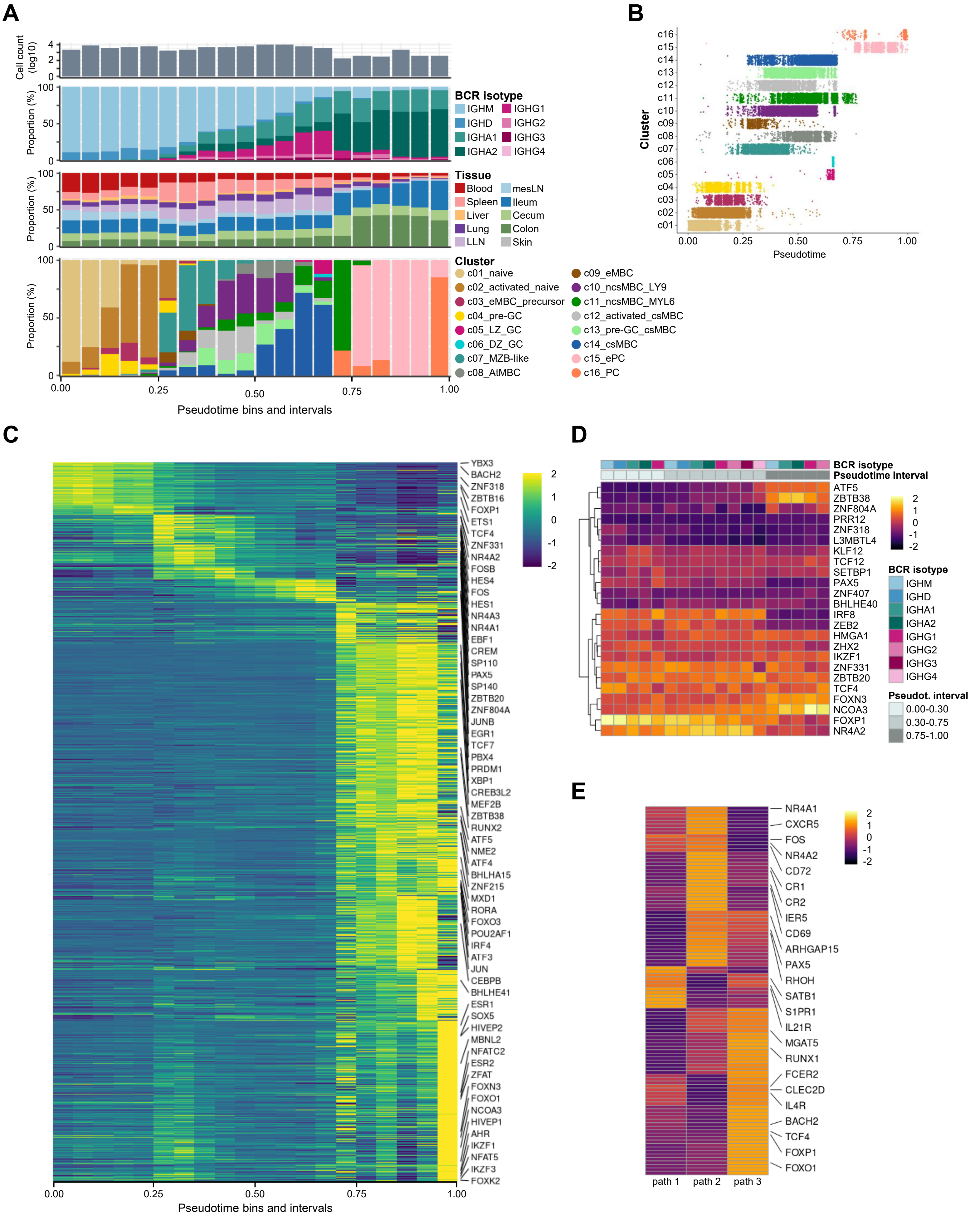

**Figure S4. Expanded analyses of B-cell transcriptional dynamics along pseudotime, related to Figure 3.** (**A**) Bar plot showing the number of cells per pseudotime bin (upper panel); and stacked bar plots illustrating isotype composition (middle-upper panel), tissue composition (middle-lower panel), and cluster composition (lower panel) for each pseudotime bin. (**B**) Stacked dot plot showing the distribution of individual B cells across pseudotime, organized and coloured by B-cell cluster. (**C**) Heatmap showing the log fold change of the top 50 differentially expressed genes across pseudotime, averaged per pseudotime bin. Transcription factors are highlighted in the heatmap. (**D**) Heatmap showing TFs differentially regulated across pseudotime intervals and BCR isotypes, identified through differential gene expression analysis (see Table S8). Conditions with fewer than 50 cells were excluded from the analysis to ensure robust statistical power. (**E**) Heatmap showing the log fold change of the top 50 differentially expressed genes averaged across the three developmental paths defined in Figure 3I.

### Figure S5. Related to Figure 4

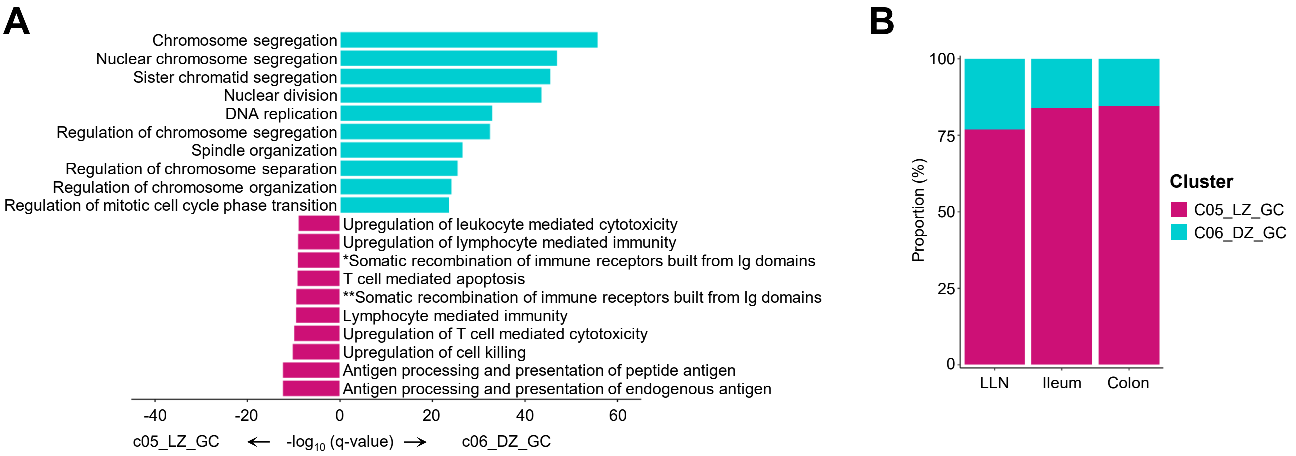

**Figure S5. Functional divergence of c05_LZ_GC and c06_DZ_GC B cell subsets across tissues, related to Figure 4.** (**A**) Bar plot showing the top Gene Ontology pathways enriched in c05_LZ_DZ and c06_DZ_GC B cell clusters compared with each other. Bar colors indicate cluster identity. Full term for *: "Positive regulation of adaptive immune response based on somatic recombination of immune receptors built from immunoglobulin superfamily domains"; full term for **: "Adaptive immune response based on somatic recombination of immune receptors built from immunoglobulin superfamily domains". (**B**) Stacked bar plot showing the tissue distribution of c05_LZ_DZ and c06_DZ_GC cells. Tissues with fewer than 50 cells in either cluster were excluded to ensure robust statistical power.

### Figure S6, related to Figure 5

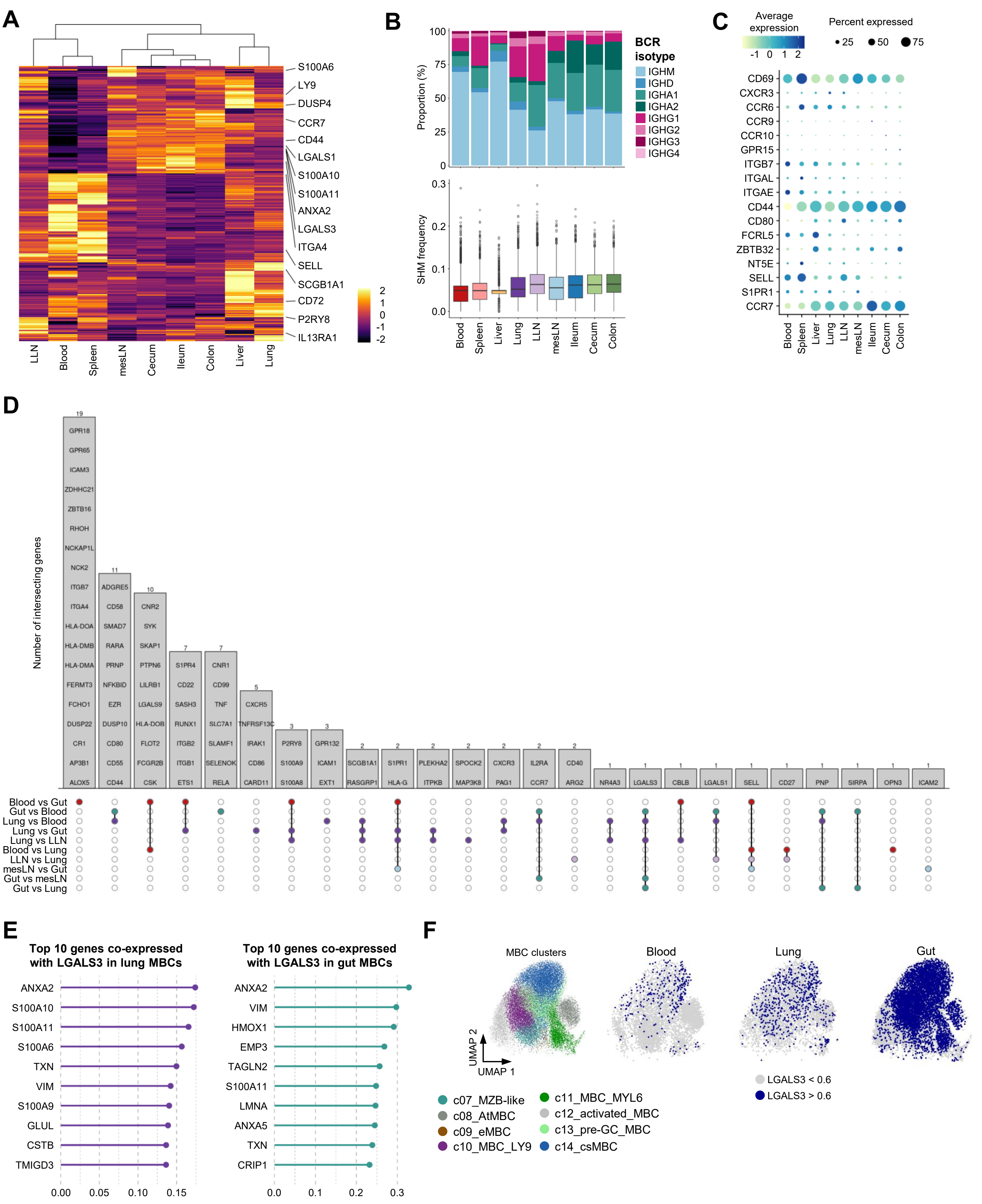

**Figure S6. Tissue-specific transcriptomic landscape of human MBCs reveals differential Ig-isotype usage and novel residency-associated markers, including an LGALS3-centered co-expression network, related to Figure 5.** (**A**) Heatmap showing the log fold change of the top 50 differentially expressed genes in MBCs averaged across tissues. (**B**) Upper panel: stacked bar plots showing the BCR isotype for each tissue for MBCs. Upper panel: Box plot of the SHM frequency on the IgH chain across tissues for MBCs. Each box represents the interquartile range (IQR) of SHM frequencies for a given cluster, while the center line represents the median. Whiskers extend to 1.5 times the IQR starting from the respective box boundary. Individual dots represent outliers. (**C**) Dot plot of the average expression of marker genes for MBCs across tissues. Dot size indicates the percentage of cells expressing each gene, and dot color represents the average gene expression level. (**D**) UpSet plot illustrating intersecting genes across the specified tissue comparisons. The analysis focused on genes related to adhesion molecules, G protein–coupled receptors (GPCRs), and extracellular matrix components (**Table S5**). Genes with expression >0.5 and a minimum twofold difference between tissues were included. (**E**) Lollipop plot showing the top 10 genes most positively co-expressed with *LGALS3* in lung (left panel) and gut (right panel) MBCs, based on Pearson correlation of log-normalized expression data. (**F**) UMAP plots of the MBC clusters colored by cell annotation (left panel), and LGALS3 expression.

### Figure S7, related to Figure 5

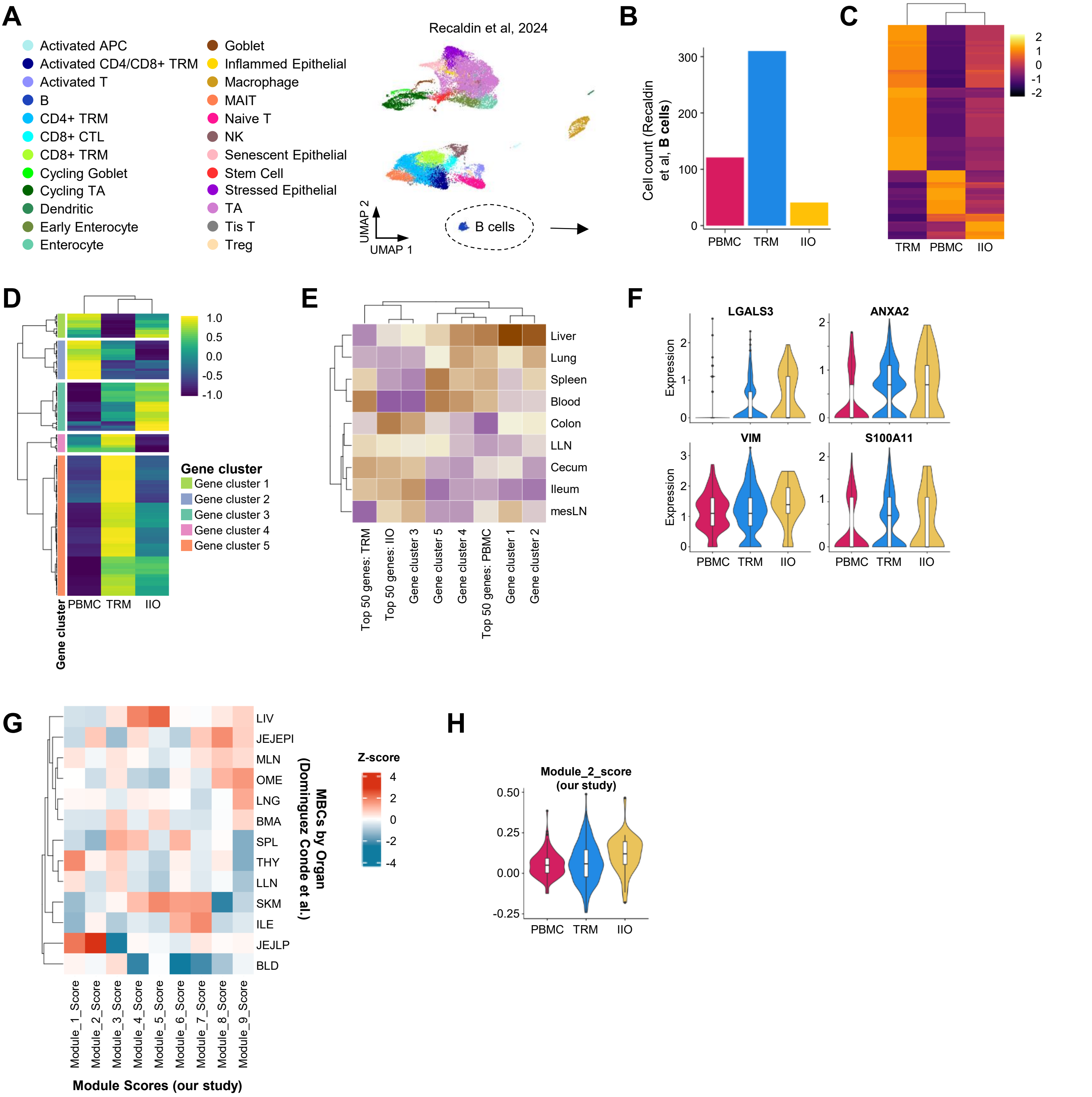

**Figure S7. Comparison of tissue-resident B-cell phenotypes with external datasets, related to Figure 5.** (**A**) UMAP projection of the Recaldin et al. dataset colored by cell clusters as originally published. (**B**) Bar plot showing the number of B cells per condition in the Recaldin et al. dataset. (**C**) Heatmap of log fold changes for the top 50 DEGs in B cells across experimental conditions from Recaldin et al. (**D**) Heatmap of log fold changes for the top 50 DEGs related to adhesion molecules, G protein–coupled receptors (GPCRs), and extracellular matrix components in B cells across conditions (Recaldin et al. dataset). (**E**) Heatmap depicting average gene signature scores derived from the DEGs in (**C**) and (**D**), projected onto MBCs from our dataset, stratified by tissue of origin. (**F**) Violin plots showing the log-normalized expression of *LGALS3* and its top co-expressed partners *ANXA2*, *VIM*, and *S100A11* across conditions (Recaldin et al. dataset). (**G**) Heatmap showing average module-specific gene signature scores (from **Figure 5A**) calculated across MBCs from the Dominguez Conde et al. dataset and grouped by organ. Conditions with fewer than 50 cells were excluded from the analysis to ensure robust statistical power. (**H**) Violin plots showing the average gene Module_2_score signature score (from **Figure 5A**) in B cells across conditions (Recaldin et al. dataset). (**F and H**) Each violin depicts the full expression distribution; the white box and central line indicate the interquartile range and median, respectively.

### Figure S8, related to Figure 6

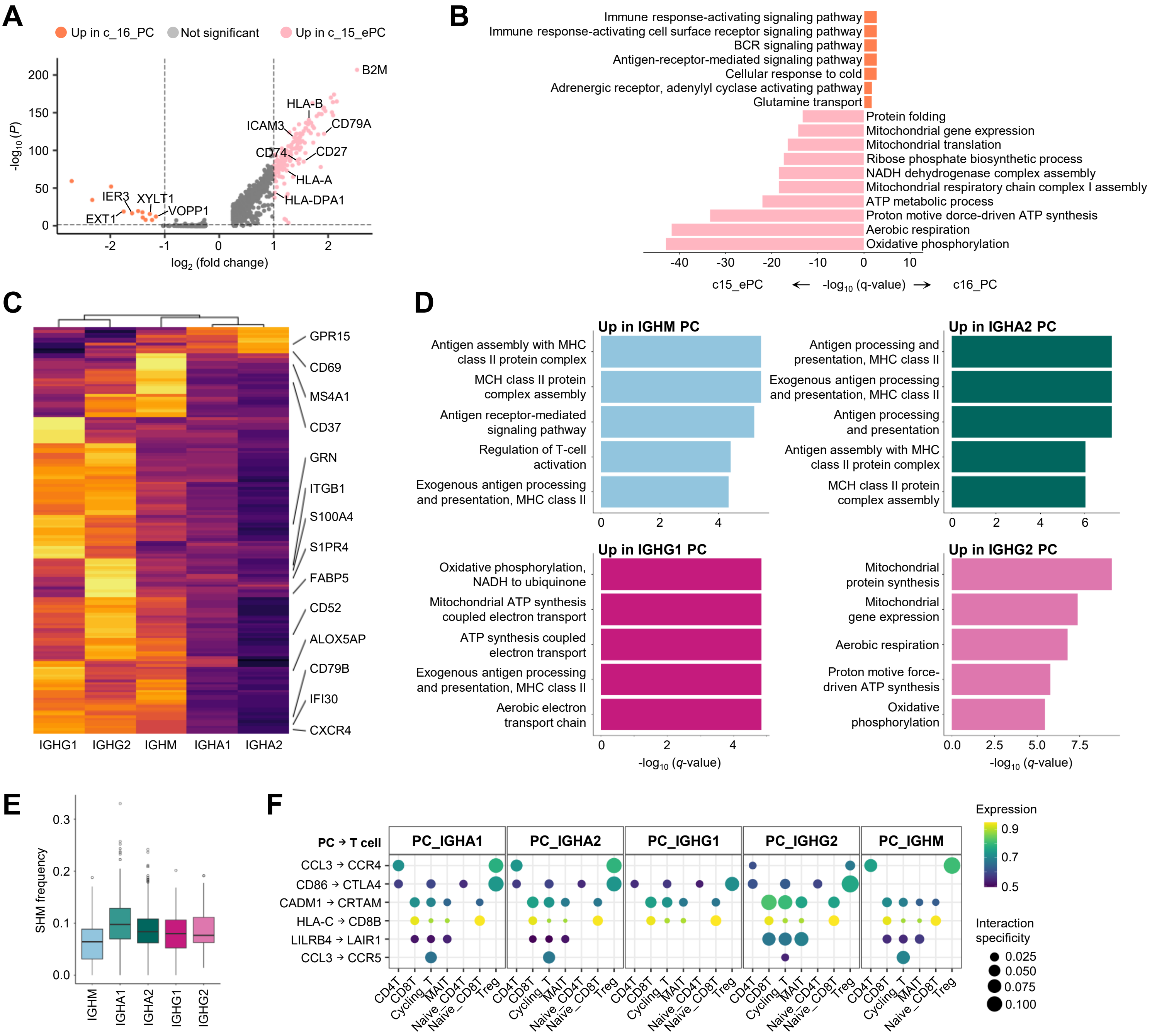

**Figure S8. Isotype-linked gene programs and functional pathways in plasma cells, related to Figure 6.** (**A**) Volcano plot showing DEGs between immature c15_ePCs and mature c16_PCs. Vertical dashed lines mark $\log_2$FC thresholds of ± 1; the horizontal dashed line indicates *FDR* = 0.05. (**B**) Bar plot showing the top Gene Ontology pathways enriched in c15_ePC and c16_PC clusters compared with each other. Bar colors indicate cluster identity. (**C**) Heatmap showing the log fold change of the top 50 differentially expressed genes in PCs averaged across isotypes. (**D**) Bar plots of top enriched Gene Ontology pathways in PCs by isotype; colors indicate BCR isotype. (**E**) Box plot of SHM frequency on the IgH chain of PCs across isotypes. Each box represents the interquartile range (IQR) of SHM frequencies for a given cluster, while the center line represents the median. Whiskers extend to 1.5 times the IQR starting from the respective box boundary. Individual dots represent outliers. (**F**) Dot plot depicting predicted ligand-receptor interactions between T cells and PCs expressing different isotypes. Dot size represents interaction specificity scores, and color intensity reflects the average expression level of the ligand-receptor pair across the interacting cell types. (**C-F**) Conditions with fewer than 50 cells were excluded from the analysis to ensure robust statistical power.
